# The HUSH epigenetic repressor complex silences PML nuclear bodies-associated HSV-1 quiescent genomes

**DOI:** 10.1101/2024.06.18.599571

**Authors:** Simon Roubille, Tristan Escure, Franceline Juillard, Armelle Corpet, Rémi Néplaz, Olivier Binda, Coline Seurre, Mathilde Gonin, Stuart Bloor, Camille Cohen, Pascale Texier, Oscar Haigh, Olivier Pascual, Yonatan Ganor, Frédérique Magdinier, Marc Labetoulle, Paul J. Lehner, Patrick Lomonte

## Abstract

Herpes simplex virus 1 (HSV-1) latently infected neurons show multiple patterns in the distribution of the viral genomes within the nucleus, at least in mouse models. One of the major patterns is characterized by the presence of quiescent HSV-1 genomes trapped in promyelocytic leukemia nuclear bodies (PML NBs) to form viral DNA-containing PML-NBs (vDCP NBs). Using a cellular model reproducing the formation of vDCP NBs we previously showed that viral genomes are chromatinized with the H3.3 histone variant modified on its lysine 9 by tri-methylation (H3.3K9me3) a chromatin mark associated with transcriptional repression. Here we identify an essential role for the HUSH complex and its SETDB1 and MORC2 effectors in the acquisition of the H3K9me3 mark on the PML NBs-associated HSV-1 and in the maintenance of HSV-1 transcriptional repression. ChiP-seq analyses highlight the association of the H3K9me3 mark with the entire viral genome. Inactivating the HUSH-SETDB1-MORC2 repressor complex prior to viral infection results in a significant reduction of H3K9me3 on the viral genome, while the overall impact on the cellular genome is minimal, except for expected changes in families of LINE1 retroelements. Depletion of HUSH, SETDB1, or MORC2, relieves the repressive state of HSV-1 in infected primary human fibroblasts as well as human induced pluripotent stem cell-derived sensory neurons (hiPSDN). We discovered that the viral protein ICP0 induces MORC2 degradation via the proteasome machinery. This process is concurrent with ICP0 and MORC2 depletion capability to reactivate silenced HSV-1 in hiPSDN. Overall, our findings underscore the robust antiviral function of the HUSH-SETDB1-MORC2 repressor complex against a herpesvirus by modulating chromatin marks linked to repression, thus presenting promising avenues for novel anti-herpesvirus therapeutic strategies.

**Significance statement:** Herpes simplex virus 1 (HSV-1) is a major human pathogen, which remains latent in the trigeminal ganglia (TG) neurons of the infected individuals. Its reactivation is characterized by a variety of clinical symptoms the most severe ones being keratitis and herpesvirus encephalitis. The colonization of the CNS by the virus during the individual life is a well-known fact but the pathophysiological effects on neurons homeostasis are still underestimated. It is thus paramount to understand the molecular mechanisms that control HSV-1 latency and maintain the virus in a pseudo silent state.

## Introduction

The establishment of latent herpes simplex virus 1 (HSV-1) infection involves a complex interplay of cellular and molecular events. At the epigenetic level, this requires the viral chromatinization and interaction of latent episomal viral genomes with their nuclear environment (1, 2). Promyelocytic leukemia nuclear bodies (PML NBs also named ND10) are nuclear membrane-less organelles involved in the transcriptional control of HSV-1 latent genomes (3, 4). The protein composition of PML NBs is highly dynamic due to their phase separation properties (5). Following HSV-1 genomes nuclear entry, PML NBs sequester HSV-1 genomes (3, 6, 7) and control the acquisition of repressive histone marks of PML NBs-associated HSV-1 genomes (also named viral DNA-containing PML NBs, vDCP NBs or ND10-like) (8). VDCP NBs, a major hallmark of HSV-1 latently infected neurons, are characterized by their ability to maintain latent HSV-1 genomes in a transcriptionally inactive, but reversible, state of quiescence (3, 4). PML NBs, and by extension vDCP NBs, are disrupted by ICP0, a virus-encoded ubiquitin E3 ligase, which counteracts the intrinsic host antiviral defense mechanisms, and is essential for the initiation of HSV-1 lytic reactivation and replication from latency (8–12). Di- and trimethylation of lysine 9 of histone H3 (H3K9me3) marks repressive heterochromatin in general and is with H3K27me3 associated with repressed latent HSV-1 genomes (1, 8, 13–15), but the cellular factor(s) involved in the establishment and maintenance of HSV-1 latency remain poorly defined (16–20). Three methyltransferases are responsible for the deposition of H3K9me3 in mammalian cells: SUV39H1/2 and ESET/SETDB1 (hereafter named SETDB1) (21–23). In mouse cells SETDB1 is a major constituent of PML NBs (24). SETDB1 is responsible for silencing proviral elements in ESC cells (25) and for the establishment and maintenance of human cytomegalovirus latency in CD34+ cells (26). We identified HUSH (Human Silencing Hub) as an epigenetic repressor complex which silences integrated retroelements (27, 28). These include retroviruses, such as latent HIV (27), as well as retrotransposons (29–33). HUSH also suppresses recombinant adeno-associated virus (rAAV) (34), and normally integrated viruses that have been engineered to remain unintegrated, such as murine leukemia virus (MLV) (32). HUSH made up of three core components, TASOR (transgene activation suppressor), MPP8 (M-phase phosphoprotein 8) and Periphilin (PPHLN1) and recruits two effector proteins: the SETDB1 methyltransferase deposits repressive H3K9me3 heterochromatin (27), and the MORC2 (microrchidia CW-type zinc-finger) ATP-dependent chromatin remodeler compacts heterochromatin (35, 36), and is frequently mutated in Charcot-Marie-Tooth disease CMT2Z patients (35–39).

Here we investigate the role of SETDB1 and the HUSH complex in the regulation of PML NBs-associated quiescent HSV-1, an unintegrated herpesvirus genome. We find that HUSH, SETDB1 and MORC2 participate to an epigenetic environment which allows the establishment and maintenance of HSV-1 repression in vDCP NBs in primary human fibroblasts as well as sensory neurons derived from human induced pluripotent stem cells (hiPSDN), the main cells supporting HSV-1 latency.

## Results

### SETDB1 is a major determinant of vDCP NBs

To understand how the HSV-1 viral genome is maintained in the quiescent state we initially infected primary human fibroblast cells (hFC) with a quiescence-prone HSV-1 (lqHSV-1) mutant (8), a model system we previously showed to reproduce the formation of vDCP NBs, as seen in trigeminal neurons of latently infected mice and human (3, 4). Proteins of the HIRA H3.3 histone chaperone complex, HIRA and UBN1 bind naked non-chromatinised DNA (40) and are recruited to vDCP NBs where they bind incoming HSV-1 genome (8). LqHSV-1 genome chromatinization with H3.3 by the HIRA complex and PML protein results in K9 trimethylation of the H3.3 histone variant (8). UBN1 co-precipitates with a histone H3K9 methyltransferase activity (41), and SETDB1 is a component of PML NBs in mouse cells (24). Therefore, we investigated whether SETDB1 could play a role in HSV-1 dependent silencing. SETDB1 was readily detected in PML NBs of non-infected (NI) and infected hFC (**Supplementary Fig. 1a**), remaining associated with PML NBs in lqHSV-1 infected cells for at least 7 days post-infection (dpi) (**Supplementary Fig. 1a**) and co-localizing with lqHSV-1 genomes in all infected cells (**Supplementary Fig. 1b**). These data suggest SETDB1 and its partner proteins could be putative molecular determinants in the epigenetic regulation of PML NBs-associated quiescent HSV-1 genomes.

### The HUSH complex and MORC2 accumulate in vDCP NBs following HSV-1 PML NBs-associated quiescent infection

Given the close functional interaction between SETDB1, the HUSH complex and MORC2, we investigated their presence in PML NBs and vDCP NBs in NI and infected cells. All proteins detected by IF-FISH, including endogenous SETDB1 and MPP8, were associated with vDCP NBs in quiescently infected hFC (**Fig. 1**) and were present in the same structures as exemplified by the concomitant detection of SETDB1 and MPP8 co-localizing with viral genomes (**Supplementary Fig. 1c**). Unlike endogenous SETDB1 and over-expressed proteins, none of the endogenous core HUSH complex proteins or MORC2 showed PML NBs localization in NI hFC (**Supplementary Fig. 2 and Supplementary Fig. 3**). We performed ChIP analysis to determine whether the HUSH complex and its effector proteins were recruited on the HSV-1 genome. Multiple loci (promoter and/or coding region, see MM for details) within the viral genome were analyzed, which together represent all classes of lytic and latent genes (**Fig. 2a**). Only the antibody against endogenous MPP8 gave reliable results by ChIP, and showed a strong interaction between MPP8 and all viral loci (**Fig. 2b, and Supplementary Fig. 5a** for interactions of endogenous MPP8 with representative cellular loci in hFC). Ectopic tagged HUSH components, including MPP8, were also enriched on the quiescent viral genome (**Fig. 2c, and expanded data** in **Supplementary Fig. 4**). We have previously demonstrated that the interaction of chromatin modulator including the H3.3 chaperone proteins HIRA, UBN1, DAXX, and ATRX across the lqHSV-1 genome is a specific feature of the quiescent state corresponding to the association of HSV-1 genomes with PML NBs (8). Together, these data suggest a role for the HUSH complex, together with SETDB1 and MORC2, in the epigenetic regulation of PML NBs-associated quiescent HSV-1 genomes.

**Fig. 1.**
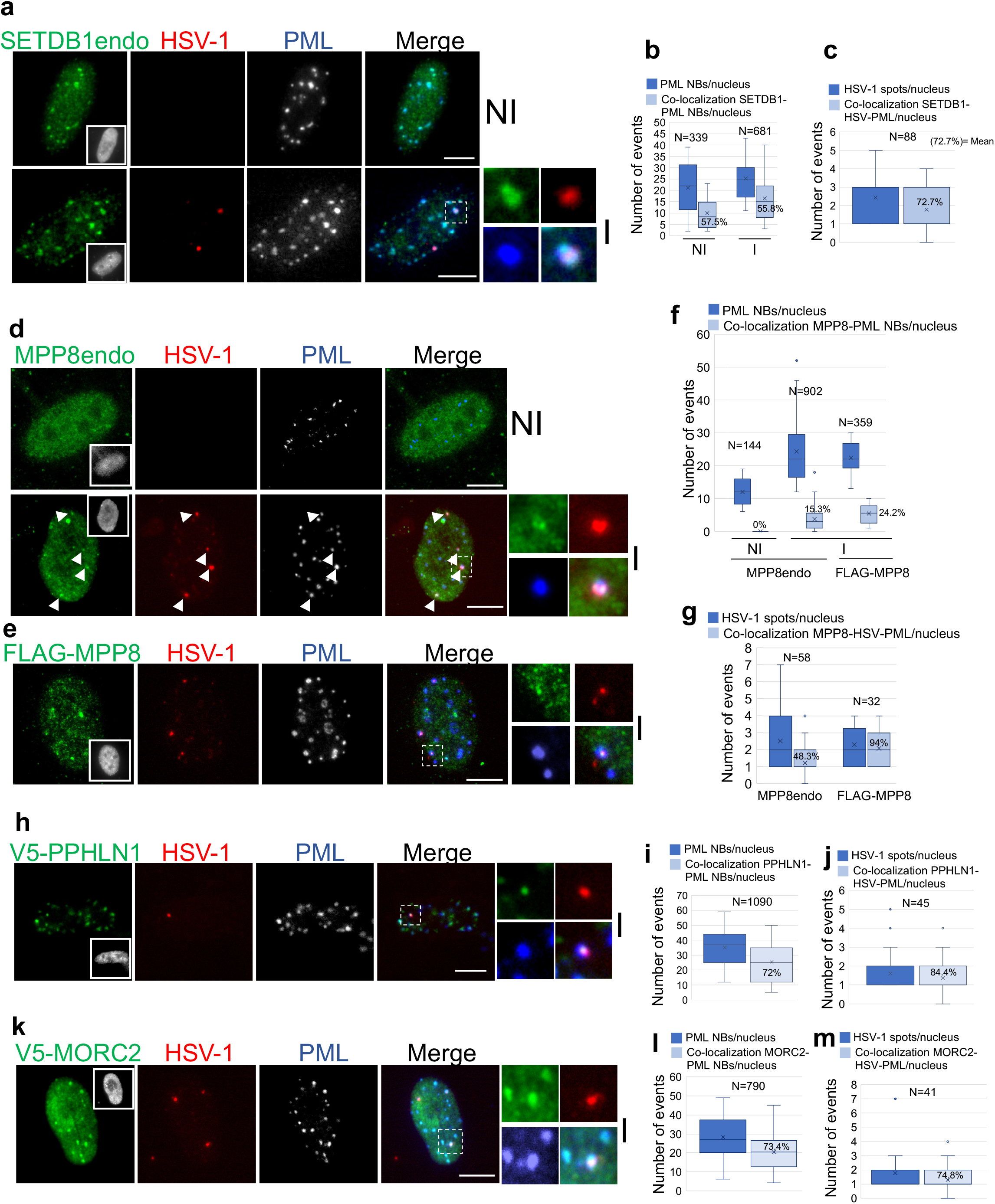
Detection of proteins of the HUSH-SETDB1-MORC2 entity in vDCP NBs. FISH-IF to detect proteins of the HUSH-SETDB1-MORC2 (green) entity and PML (blue/grey) together with HSH-1 genomes. hFC were infected or not with lqHSV-1 (m.o.i.= 3, 100% of cells infected) and harvested at 4 dpi to perform immuno-FISH. Endogenous SETDB1 and MPP8 were readily detectable with native antibodies supporting the FISH procedure. All other proteins were detected through the expression of an ectopic tagged version. FLAG-MPP8 was also detected as a control for the other HUSH complex components. Insets represent nuclei. Zooms point out vDCP NBs. **a, d, e, h, k,** detection of endogenous SETDB1, endogenous MPP8, FLAG tagged MPP8, v5 tagged periphilin (PPHLN1), and v5 tagged MORC2, respectively. NI = not infected; I = infected. Scale bars represent 5µm. **b, f, i, l,** quantification of the average number of PML NBs per nucleus (dark blue) and of the co-localization of the protein in the PML NBs (light blue) for the respective samples in NI and/or I cells. Means are indicated in the graphs. **c, g, j, m,** quantification of the average number of HSV-1 spots per nucleus (dark blue) and of the co-localization of the protein with HSV-1 and PML (light blue) for the respective samples in infected cells. N = number of events counted. Means are indicated in the graphs.

**Fig. 2.**
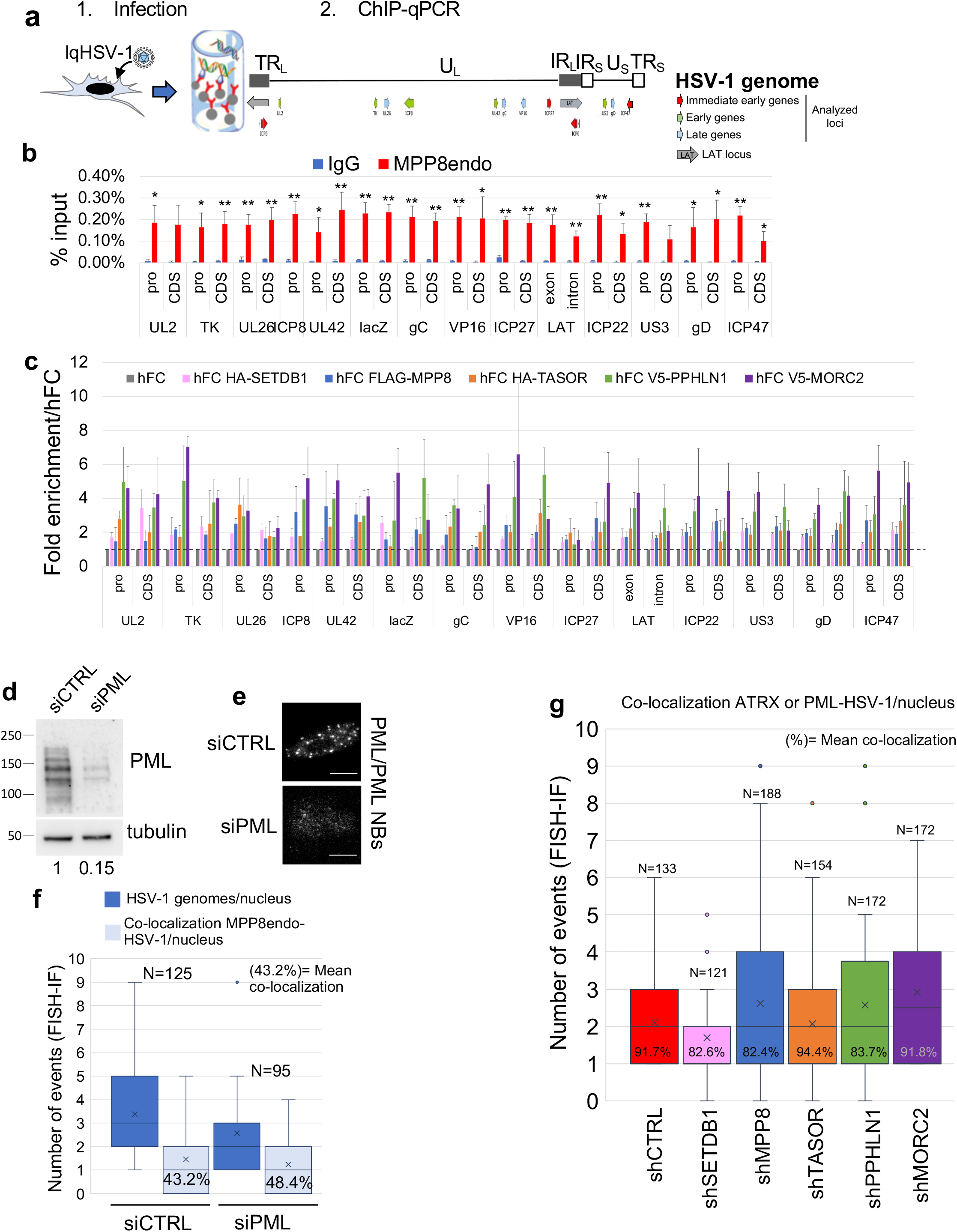
Interaction of the proteins of the HUSH-SETDB1-MORC2 entity with PML NBs-associated quiescent HSV-1 genome. **a,** schematic view of the ChIP procedure to detect interactions of proteins with viral genomes. Control hFC or hFC expressing a tagged version of proteins of interest were infected with lqHSV-1 (m.o.i.= 3) and ChIP-qPCR was performed 24h pi. All experiments were performed at least 3 times. Measurements were taken from distinct samples. Quantifications data are mean values (+/- SD). P-values * <0.05; ** <0.01 (one-tailed paired Student’s t-test). **b,** interaction of endogenous MPP8 with viral genome loci. **c,** mean fold enrichments on the different viral loci of the corresponding proteins compared to control hFC. For quantification details on each loci see expended **Supplementary Fig. 4. d-g**, MPP8endo co-localization with lqHSV-1 in hFC transfected with a siRNA CTRL (siCTRL) or siRNA against PML (siPML) and infected with lqHSV-1 for 48h (m.o.i.= 3). **d**, WB of PML in siCTRL or siPML-treated hFC. The quantification of PML levels relative to tubuline is depicted below the WB. **e**, Representative images showing the PML NBs signal by immunofluorescence (grey) to exemplify the efficiency of the siPML on PML NBs disappearance (also see (85)). **f**, Quantification of the co-localization of MPP8endo with lqHSV-1 following immuno-FISH to detect MPP8, lqHSV-1, and PML. Distribution of HSV-1 genome signals/nucleus (dark blue) and co-localization of MPP8endo with lqHSV-1/nucleus (light blue) are shown. Means of colocalization between MPP8endo and lqHSV-1 are shown (in %). N= total number of viral genomes analyzed. **g**, Quantification of ATRX co-localization with lqHSV-1 in hFC transduced with lentiviruses expressing control shRNA (shCTRL) or shRNA against proteins of interest and infected with lqHSV-1 for 48h (m.o.i.= 3). Distribution of the numbers of co-localization between ATRX and lqHSV-1/nucleus are shown. Means of colocalization between ATRX and lqHSV-1 are shown (in %). N= total number of viral genomes analyzed.

### PML and the HUSH-SETDB1-MORC2 repressor complex are independent for their interaction with lqHSV-1

In order to investigate the interaction and/or co-localization of HSV-1 genomes with HUSH-SETDB1-MORC2 proteins in the absence of PML, PML-depleted hFC (8) were infected with lqHSV-1, and ChIP-qPCR or FISH-IF were conducted. The ChIP results show that depleting PML before infection does not affect the interaction between endogenous MPP8 with the lqHSV-1 genomes (**Supplementary Fig. 5b-d**). Quantification of FISH-IF data reveals that the depletion of PML does not decrease the co-localization of endogenous MPP8 with lqHSV-1 genomes (**Fig. 2d-f)**. Similarly, individual depletion of HUSH-SETDB1-MORC2 proteins does not impact on the formation of vDCP NBs as monitored by the co-localization of PML or ATRX, one of the major components of PML NBs, with lqHSV-1 (**Fig. 2g, and Supplementary Fig. 6**). These findings suggest that, while ultimately all cellular components are present in the vDCP NBs, the interaction between lqHSV-1 genomes with PML/PML NBs and the HUSH-SETDB1-MORC2 repressor complex may occur independently of each other.

### The HUSH complex with SETDB1 and MORC2 control repressive histone marks on quiescent PML NBs-associated HSV-1 genome

To further investigate the role of HUSH components in the acquisition of the H3K9me3 mark on the lqHSV-1 genome, we took a knock down approach, transducing hFC with previously validated and specific shRNAs (27, 35) (**Fig 3a**) and verified their efficiency at the protein level (**Fig. 3b-f**). With the unexpected exception of TASOR, depletion of the proteins induced a significant decrease of the H3K9me3 mark across the quiescent viral genome (**Fig. 3g and expanded data in Supplementary Fig. 7**). A siRNA targeting SETDB1 gave similar results (**Supplementary Fig. 8**). The reduction in H3K9me3 levels on the lqHSV-1 genome could not be linked to a general decrease of the total H3K9me3 mark in the shRNA-treated cells, as observed in Western blotting (**Supplementary Fig. 9a**). Moreover, the deficit in H3K9me3 on the lqHSV-1 genome did not stem from a general decrease in histone H3 binding, as indicated by ChIP analyses (**Supplementary Fig. 9b-f**). In contrast to SETDB1, SUV39H1, another H3K9me3 methyltransferase (22), was not detectable in the PML NBs in both non-infected and infected hFC (**Supplementary Fig. 10a**). Depletion of SUV39H1 via specific siRNA did not diminish the levels of H3K9me3 associated with the lqHSV-1 genome (**Supplementary Fig. 10b-e**).

**Fig. 3.**
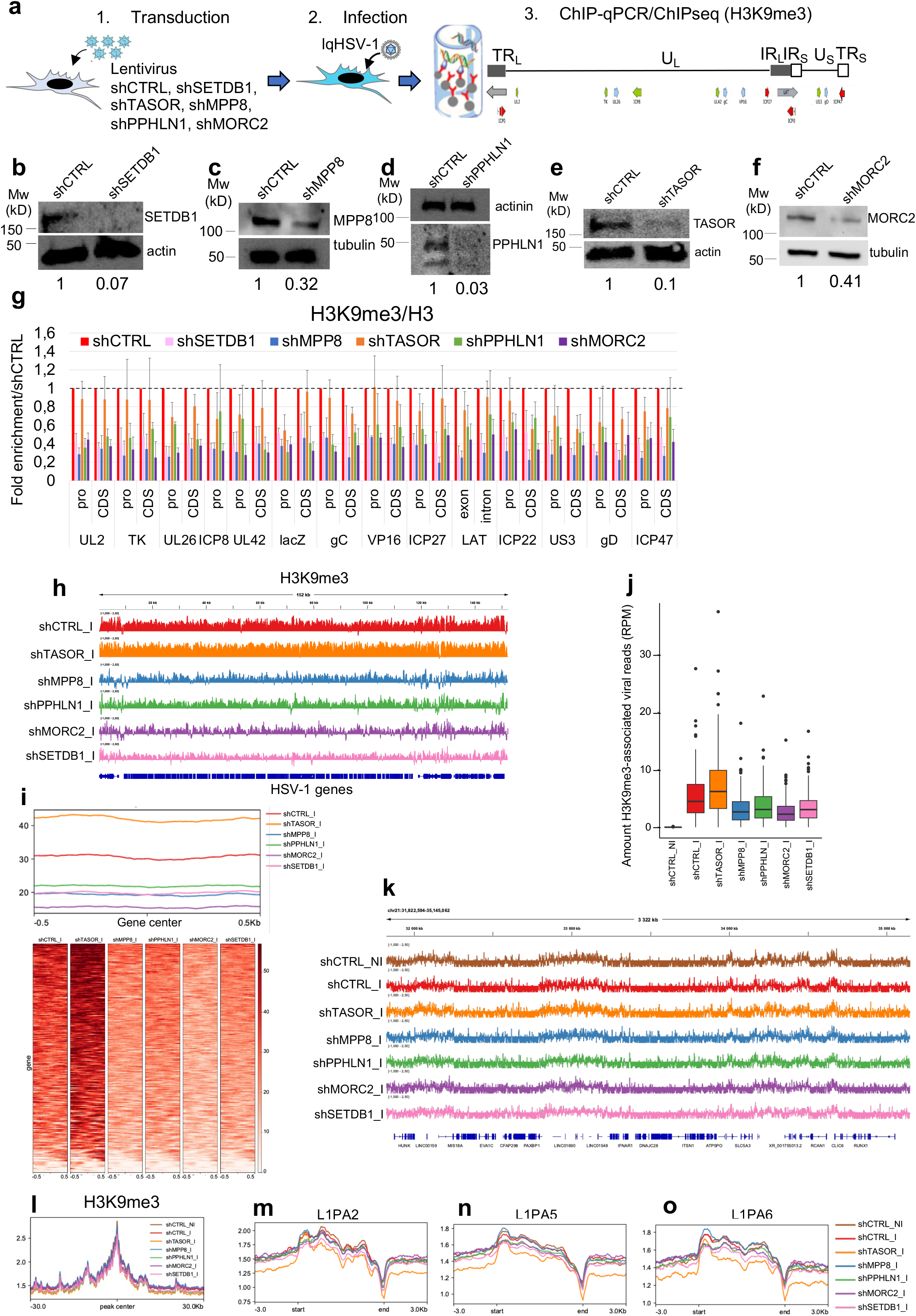
Inactivation of proteins of the HUSH-SETDB1-MORC2 entity reduces H3K9me3 mark on PML NBs-associated quiescent HSV-1 genome. **a,** schematic view of the ChIP-qPCR/ChIP-seq procedure to detect enrichment of H3K9me3 on viral genomes after depletion or not by shRNAs of proteins of interest. **b-f,** WB on the target proteins to confirm the efficiency of the specific shRNAs. The quantification of protein levels relative to housekeeping proteins is depicted below the WBs. **g,** ChIP-qPCR showing mean fold enrichments on the different viral loci of H3K9me3 in specific shRNA-depleted cells compared to cells expressing a control shRNA. Experiments were performed at least 3 times. H3K9me3 signal was standardized over total H3 present on each locus. Measurements were taken from distinct samples. Quantifications data are mean values (+/- SD). For quantification details on each loci see expanded **Supplementary Figure 7**. P-values * <0.05; ** <0.01; *** <0.001 (one-tailed paired Student’s t-test). **h to o,** data from H3K9me3 ChIP-seq experiments in infected hFC treated with shCTRL or a shRNA against the indicated proteins of interest. **h,** Genome browser snapshot of the H3K9me3 enrichment normalized on input across the entire quiescent HSV-1 genome. **i,** Profile plot and heatmaps showing the density of H3K9me3 signal on HSV-1 genes. **j,** Box plots showing the quantification of H3K9me3 reads on HSV-1 genes (in Reads Per Million). **k,** Representative genome browser snapshot of the H3K9me3 enrichment normalized on input across a gene-rich region of chromosome 21. **l,** Profile plot of the H3K9me3 density on large H3K9me3-rich regions defined in (42). **m, n, o,** Profile plots of the H3K9me3 density on specific families of LINEs regulated by HUSH.

Next, we conducted ChIP sequencing (ChIP-seq) in infected cells to comprehensively examine the deposition of the H3K9me3 mark on both the lqHSV-1 and cellular genomes. Our findings confirmed the presence of this mark across the entire viral genome and demonstrated its dependence on the activity of the HUSH-SETDB1-MORC2 repressor complex (**Fig. 3h and i, and supplementary Fig. 11a).** As previously observed through ChIP-qPCR, depletion of TASOR led unexpectedly to a general increase in the H3K9me3 mark on the viral genome. Quantitative analysis of the viral reads associated with H3K9me3 across different conditions confirmed that, except for the shTASOR-treated cells, which showed a clear increase, the number of viral reads detected in infected cells treated with HUSH-SETDB1-MORC2 shRNAs was lower compared to control shRNA-treated cells **(Fig. 3j).** No viral reads were counted from uninfected cells **(Fig. 3j).** The overall reduction in the H3K9me3 mark on the lqHSV-1 genome did not stem from a general effect of the inactivation of the HUSH-SETDB1-MORC2 repressor complex, as this decrease was not universally observed across the cellular genome **(Fig. 3k**, chr21 gene-rich region as an example), and on previously identified H3K9me3-rich large heterochromatin regions **(Fig. 3l, and Supplementary Fig. 11b)** (42). As expected, peak annotation revealed that the majority of H3K9me3 peaks was linked with non-coding regions (distal intergenic and introns) **(Supplementary Fig. 11c and d)**. Subsequently, our focus shifted towards families of long interspersed nuclear elements (LINEs) retrotransposons, which have been previously identified as undergoing significant regulation by components of the HUSH complex (27, 29, 30, 43, 44). As anticipated, the depletion of components within the HUSH-SETDB1-MORC2 complex, particularly TASOR, significantly affected the deposition of H3K9me3 on the loci **(Fig. 3m, n, o**, and quantifications of the signal on families of LINEs in **Supplementary Fig 11e**).

These findings strengthen the role of the HUSH complex, alongside SETDB1 and MORC2, as a key pathway in acquiring the repressive chromatin H3K9me3 mark on the PML NB-associated quiescent HSV-1 genome. Additionally, they suggest that TASOR inactivation may exert varying modulatory effects on H3K9me3 deposition depending on the integrated or episomal nature of specific genome loci.

### HUSH-SETDB1-MORC2 entity maintains PML NBs-associated HSV-1 quiescence

LqHSV-1 is thermosensitive and acquires its quiescent state at 38.5°C in hFC, leading to the formation of vDCP NBs and transcriptionally silent viral genomes. Latency can be reversed by destabilizing vDCP NBs with the HSV-1 protein ICP0 and reverting the temperature to 32°C (8). To determine if the HUSH complex is required to maintain lqHSV-1 silencing, individual components of the HUSH-SETDB1-MORC2 complex were depleted after the acquisition of the quiescent state and formation of the vDCP NBs (**Fig. 4a**). Inducible shRNA-mediated depletion (**Supplementary Fig. 12a-e**) of HUSH complex components led to transcriptional activation of the lqHSV-1, as determined by the appearance of viral lytic gene-encoding mRNAs (**Fig. 4b-e**). Additionally, within individual viral plaques visible in the cell monolayers displaying signs of reactivation (**Fig. 4f**), distinctive viral replication compartments (RC) indicative of ongoing viral lytic infections were easily identifiable. Notably, viral plaque formation was observed on hFC monolayers subsequent to HUSH depletion, contrasting the absence of plaque formation in the control (shCTRL) condition (**Supplementary Fig. 12f**). To confirm that inactivation of the HUSH complex induced production of infectious viral progeny, supernatants from the hFC monolayers were titrated on human bone osteosarcoma (U2OS) epithelial cells (**Fig. 4g up**). Supernatants from HUSH-depleted, but not shCTRL components induced viral plaque formation (**Fig. 4g**, supernatants of shCTRL and shMORC2 (shown as an example) and **Fig. 4h** for quantification). The HIV-2 and SIV Vpx accessory protein degrades TASOR and destabilizes the HUSH complex (31, 45). Expression of HIV-2_ROD_ Vpx also induced transcriptional reactivation, appearance of RCs, and viral plaques (**Supplementary Fig. 13a-d**) and production of infectious virus (**Supplementary Fig. 13e-f**). However, presently, we cannot dismiss the possibility of additional effects of Vpx on the stability of other cellular targets, such as SAMHD1 (46, 47), or TASOR2 a paralog of TASOR recently described as being part of a HUSH2 complex (48). These data demonstrate that the HUSH epigenetic repressor complex, along with SETDB1 and MORC2, is required for the stabilization of vDCP NBs and the PML NB-associated quiescence of HSV-1.

**Fig. 4.**
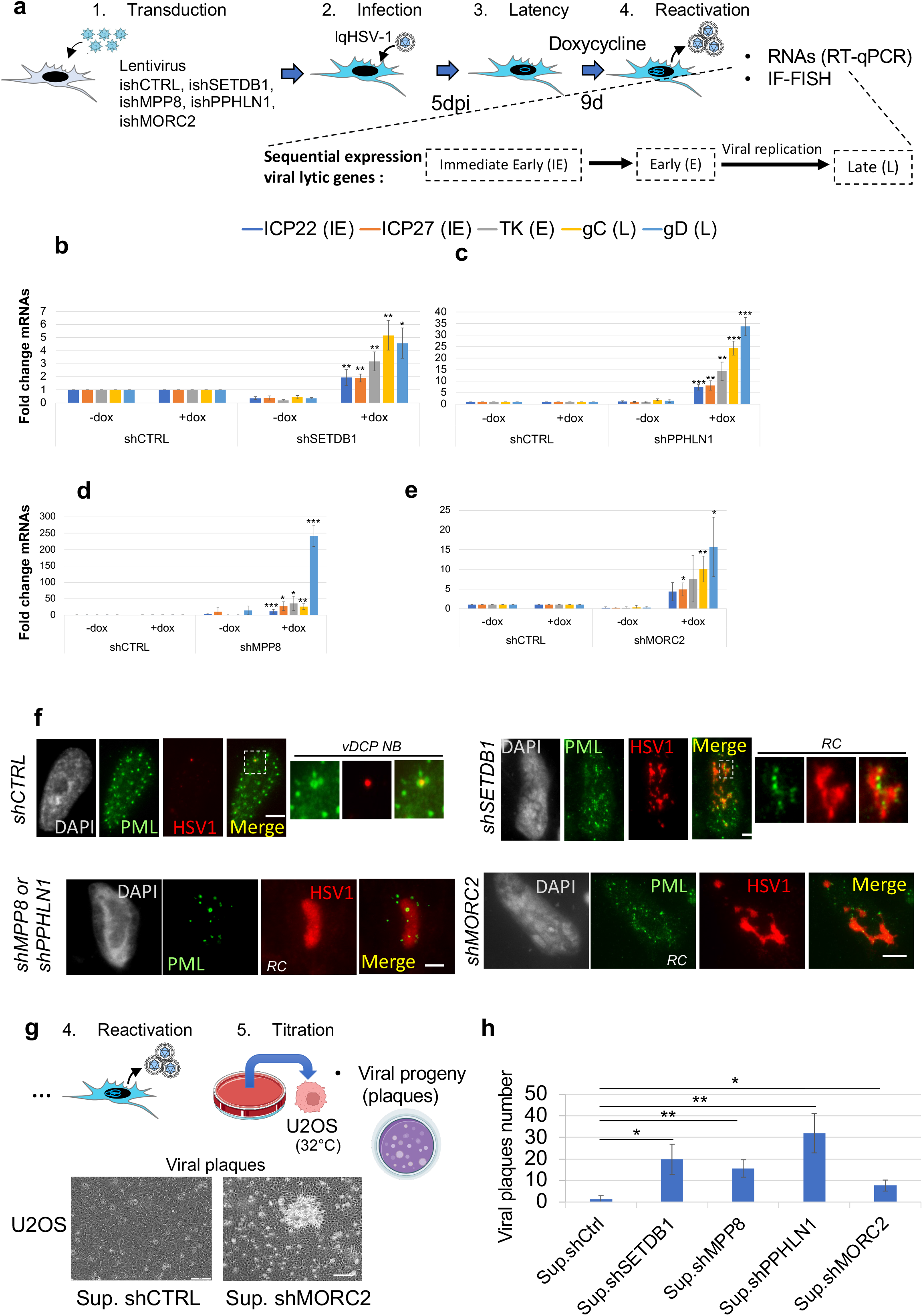
The HUSH-SETDB1-MORC2 entity maintains the silencing of PML NBs-associated quiescent HSV-1. **a,** schematic view of the experimental procedure to determine the restrictive activity of the individual proteins of the HUSH-SETDB1-MORC2 entity. hFC expressing inducible shRNA control (shCTRL) or shRNA against a protein of interest were infected with lqHSV-1 (m.o.i.= 3) at 38.5°C to establish quiescence. Five days later doxycycline was added for 9 days shifting the temperature at 32°C to allow virus replication. Cells were used to perform RT-qPCR to quantify expression of viral lytic genes or immuno-FISH to detect viral RC. **b-f,** RT-qPCR on representative HSV-1 lytic genes following treatment with a shCTRL, or a shRNA inactivating SETDB1, PPHLN1, MPP8, MORC2, respectively. All experiments were performed at least 3 times. Measurements were taken from distinct samples. Quantifications data are mean values (+/- SD). P-values * <0.05; ** <0.01; *** <0.001 (one-tailed paired Student’s t-test). **g**, immuno-FISH allowing the detection of PML (green), HSV-1 (red) and nuclei (grey, DAPI) in cell monolayers after induction of the shRNA CTRL, or targeting HUSH (shMPP8 shown as an example), or MORC2. Scale bar represents 5 µm. RC : replication compartment. **h,** titration on U2OS cells of infectious viral progeny production from above experiments using shRNA CTRL, or a shRNA inactivating each of the protein of the HUSH-SETDB1-MORC2 entity. Up : procedure; down : viral plaques with supernatant from shMORC2 hFC shown as an example. Scale bar represents 100 µm. **i**, numerations of viral plaques from the above conditions.

### The viral protein ICP0 induces the proteasome-dependent degradation of MORC2

ICP0, a RING finger-containing E3 ubiquitin ligase, is recognized for its ability to trigger the proteasomal-dependent degradation of various cellular proteins, including PML, SP100, and MORC3 (49, 50). We examined the stability of HUSH-SETDB1-MORC2 proteins in cells infected with HSV-1WT or an ICP0-deficient virus (HSV-1ΔICP0 also known as dl1403) in the presence or absence of the proteasome inhibitor MG132. As anticipated, PML isoforms were degraded in an ICP0- and proteasome-dependent manner (**Fig. 5a**). The core proteins of the HUSH complex as well as SETDB1 remained unaffected in stability upon infection. Notably, MORC2 exhibited ICP0- and proteasome-dependent degradation (**Fig. 5b**). No significant effect on MORC2 mRNA abundance dependent on expression or not of ICP0 in infected cells could be detected (**Supplementary Fig. 14**). These data establish MORC2 as a potential critical regulator in maintaining HSV-1 quiescence in neurons.

**Fig 5.**
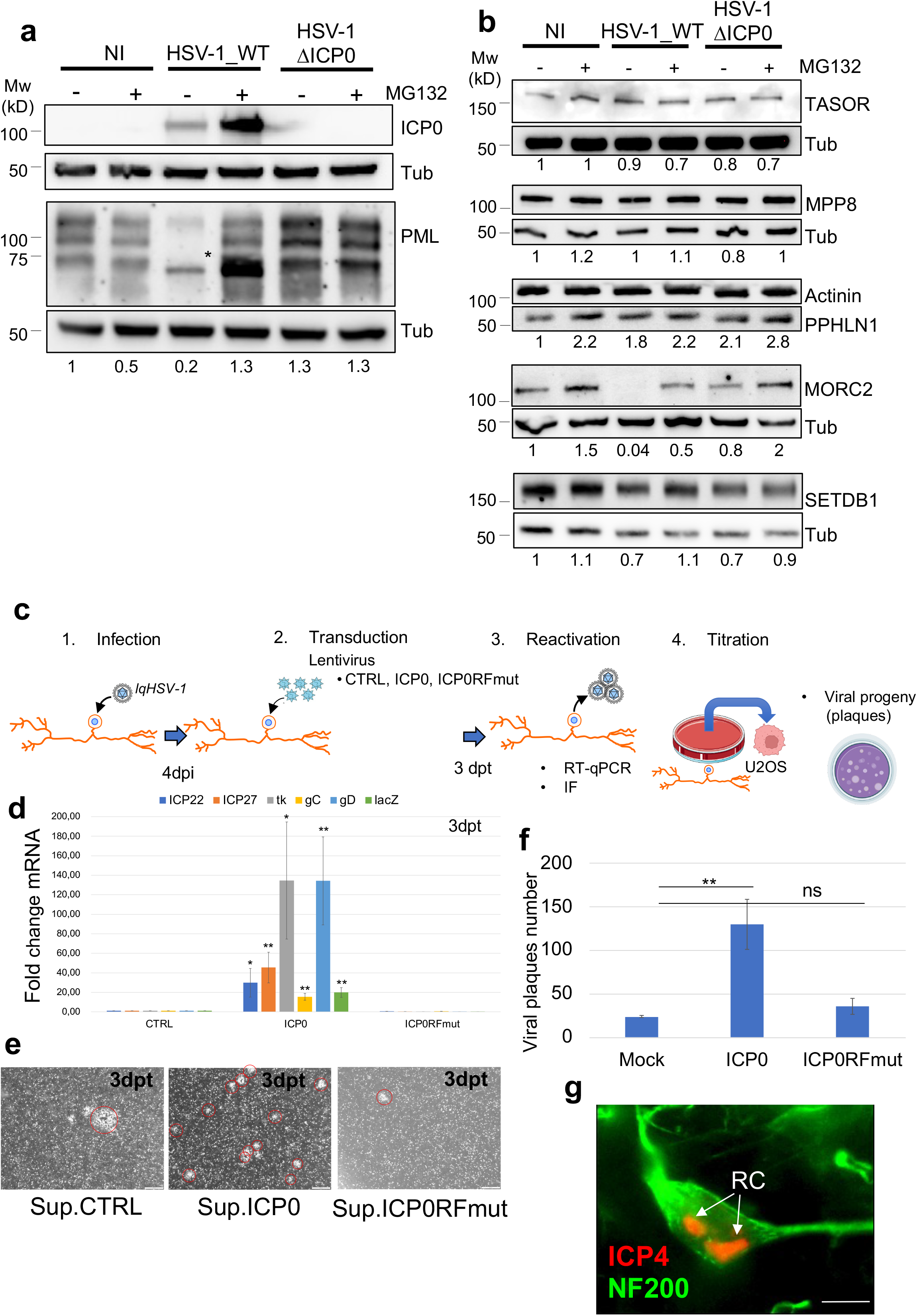
ICP0 induces proteasomal-dependent degradation of MORC2 and reactivation of HSV-1 from quiescently infected neurons. **a, b,** hFC were infected or not with wild type HSV-1 (HSV) or a mutant HSV-1 unable to express ICP0 (HSVΔICP0), in the presence (+) or not (-) of the proteasome inhibitor MG132. **a**, Detection of ICP0 expression and of PML. The star depicts the ICP0- and proteasome-dependent degradation of the SUMOylated forms of PML as first shown in (86). **b**, detection of proteins of the HUSH complex, MORC2 and SETDB1. Tubulin (Tub) or actinin were used as loading controls. Data are representative of at least 2 independent experiments that showed similar results. The quantification of protein levels relative to housekeeping proteins is depicted below the WBs. **c,** schematic view of the experimental procedure in hiPSDN infected with lqHSV-1 at 38.5°C. Four days after infection with lqHSV-1 neurons were transduced with lentiviruses control (CTRL) or expressing ICP0 or ICP0RFmut, and temperature was shifted to 32°C to allow virus replication. Multiple analyses were performed at 3 days post transduction (dpt). **d,** RT-qPCR on representative HSV-1 lytic genes following expression of ICP0 (left) or its non-functional mutant ICP0RFmut (right). All experiments are in triplicates. All quantifications data are mean values (+/- SD). Measurements were taken from distinct samples. P-values * <0.05; ** <0.01; *** <0.001 (one-tailed paired Student’s t-test). **e,** Detection of viral plaques on U2OS cell monolayers in contact with supernatants from lqHSV-1 infected neurons transduced for 3dpt with lentivirus CTRL or expressing ICP0 or ICP0RFmut. Crystal violet (grayscale) images for plaques (circled in red) visualization are shown. Scale bar represents 500 µm. **f**, numeration of the plaques detected in **e**. **g,** Immunofluorescent detection of ICP4 (red) and NF200 (green) showing viral RC in neurons. Arrows point out two RCs in the same nucleus of a single neuron. Scale bar represents 10 µm.

### MORC2 maintains PML NBs-associated HSV-1 quiescence in human neurons

The hFC infection model provides a practical system to characterize essential epigenetic features controlling PML NB-associated HSV-1 quiescence. As sensory neurons of the trigeminal ganglia (TG) are the predominant sites of latent HSV-1 infection including the formation of PML NBs-associated quiescent HSV-1, it was essential to determine whether HUSH is also able to restrict HSV-1 infection in these cells. We therefore established PML NBs-associated quiescent HSV-1 infection in human sensory neurons derived from human induced pluripotent stem cells (hiPSDN) **(Supplementary Fig. 15a)**. These neurons recapitulate the molecular, electrophysiological, and biochemical features of sensory neurons in general, and nociceptive neurons in particular **(Supplementary Fig. 15b-g)**.

HiPSDN exhibit PML NBs (**Supplementary Fig. 15h**), mirroring the presence observed in murine and human TG neurons (3, 4), and demonstrate the formation of vDCP NBs (**Supplementary Fig. 15i**). To confirm the validity of the hiPSDN infection model, we infected hiPSDN with the lqHSV-1 virus and measured reactivation following expression of the ICP0 viral protein or its inactive RING Finger domain mutant (ICP0RFmut) (**Fig. 5c**). ICP0, but not its RF mutant, destabilizes the PML NBs and vDCP NBs (8, 9) and induced transcriptional reactivation of lqHSV-1 (**Fig. 5d**), resulting in production of infectious viral progeny (**Fig. 5e-f**). Immunofluorescent detection of the ICP4 and NF200 proteins further confirmed the presence of RCs in reactivating neurons (**Fig. 5g**). Finally, shRNA-mediated MORC2 depletion from lqHSV-1-infected hiPSDNs (**Fig. 6a, and Supplementary Fig. 16**) also induced the transcriptional reactivation of lqHSV-1, as determined by the temporal increase in lytic viral mRNAs (**Fig. 6b**), confirming a role for the HUSH complex in the transcriptional silencing of PML NBs-associated quiescent HSV-1. Furthermore, infectious progeny were released from reactivating neurons, as supernatants from day 7 shMORC2-treated neurons induced viral plaques in U2OS cells (**Fig. 6c-d**). Taken together, these data demonstrate the essential role of MORC2, and by extension the HUSH complex, in maintaining PML-NBs-associated HSV-1 quiescence in neurons.

**Fig. 6.**
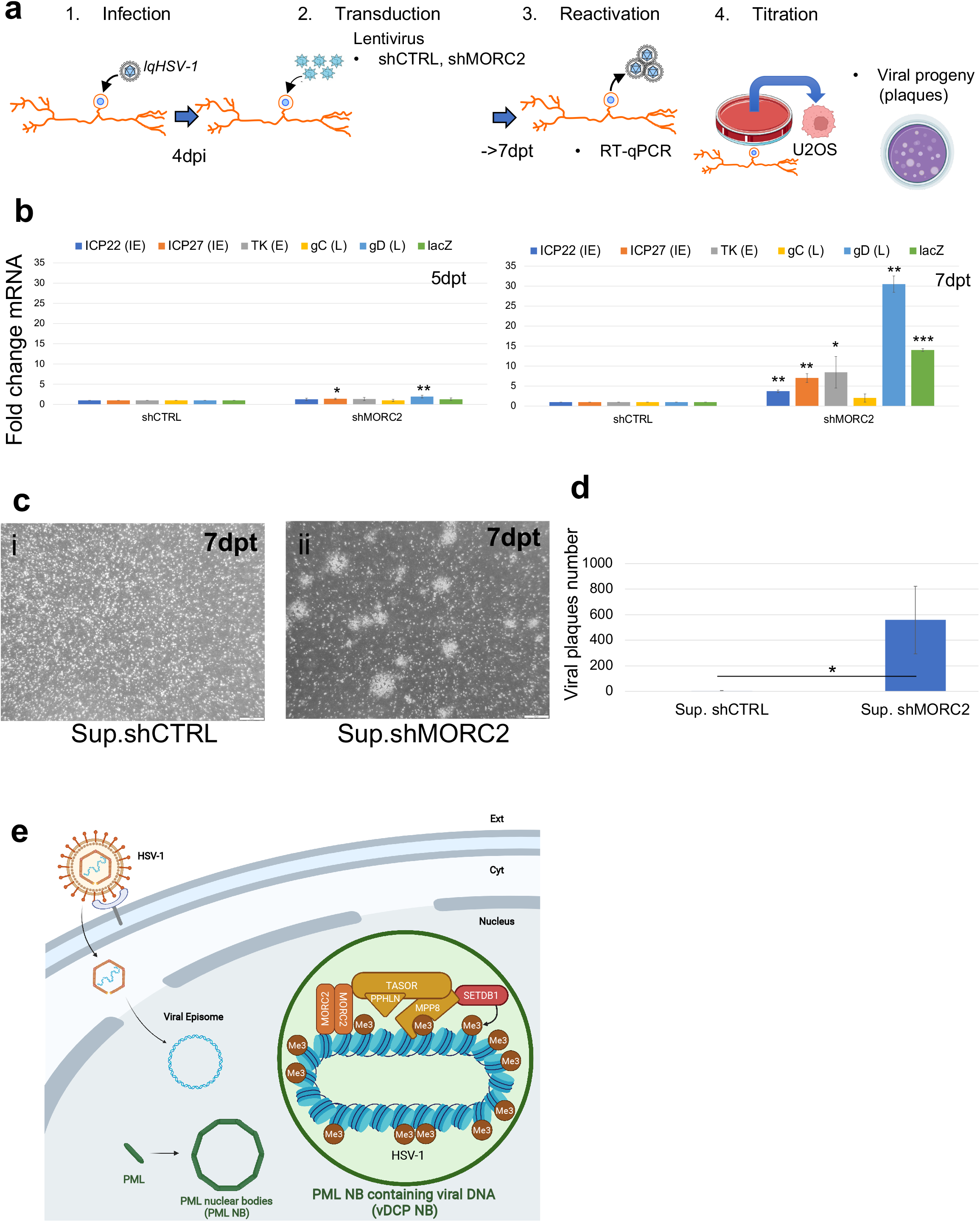
The HUSH-SETDB1-MORC2 entity acts as a restriction pathway in neurons infected with HSV-1. **a,** schematic view of the experimental procedure to determine the restriction activity of the HUSH-SETDB1-MORC2 entity in hiPSDN infected with lqHSV-1 at 38.5°C. Four days after infection with lqHSV-1 neurons were transduced with lentiviruses expressing a shRNA control, or a shRNA against MORC2, and temperature was shifted to 32°C to allow virus replication. Multiple analyses were performed 5 and 7 days post transduction (dpt). **b,** RT-qPCR at 5dpt (left) and 7dpt (right) on representative HSV-1 lytic genes following treatment with a shRNA CTRL (shCTRL), or a shRNA inactivating MORC2. All experiments are in triplicates. Measurements were taken from distinct samples. All quantifications data are mean values (+/- SD). P-values * <0.05; ** <0.01 (one-tailed paired Student’s t-test). **c,** Detection of viral plaques in U2OS cells monolayers in contact with supernatants from shCTRL (i) or shMORC2 (ii) treated lqHSV-1 infected neurons. Crystal violet (grayscale) images for plaques visualization are shown. Scale bar represents 100 µm. **d,** Numeration of viral plaques from **c**. P-values * <0.05 (one-tailed paired Student’s t-test). Measurements were taken from distinct samples. Crystal violet images (grayscale) of U2OS cell monolayers are shown above each condition. Scale bar represents 500 µm. **e,** model of epigenetic restriction of nuclear incoming HSV-1 DNA by the HUSH-SETDB1-MORC2 entity through the apposition of tri-methyl marks on H3K9 on the chromatinized viral genome ultimately leading to PML NBs-associated quiescent HSV-1 genomes. Made with Biorender.

## Discussion

The neurotropic nature of HSV-1 and its ability to spread from the peripheral to the central nervous system makes it an important etiological candidate for non-genetic infection-associated neurodegenerative pathologies (51, 52). It is therefore important to understand the mechanisms that induce and maintain HSV-1 to induce and maintain the latent state. The formation of repressive heterochromatin through K9me3 modification of histone H3.1 and/or histone variant H3.3 and chromatin compaction is an important mechanism for transcriptional silencing of endogenous retrotransposons and human immunodeficiency virus as well as HSV-1 genomes, leading to the establishment of a latent/quiescent state (8, 13, 25, 27, 28, 53, 54). Epigenetic regulation associated with viral chromatinization is therefore a hallmark of latent HSV-1. The HUSH-SETDB1-MORC2 repression complex is well described for its essential role in maintaining integrated retroelements in an inactive state (27, 35). MPP8 and SETDB1 interact and cooperate in the silencing of satellite DNA repeats in mouse embryonic stem cells (33). The HUSH complex, together with NP220/ZNF638 also silence non-integrated murine leukemia virus (32). Similarly, NP220 and HUSH silence recombinant AAV (rAAV) vectors in a serotype-dependent manner (34). However, the actual contribution of HUSH in maintaining latency of a genuine unintegrated episomal virus such as HSV-1 has not been determined.

We and others have demonstrated the heterogeneity of HSV-1 latency in neurons *in vivo* in mouse models. This heterogeneity exists at least at the level of viral genome chromatin marks acquisition and interaction of latent viral genomes with the nuclear environment. The latter is exemplified by the encapsulation of the viral genomes in a quiescent state within PML NBs, and referred as to vDCP NBs (3, 4, 13, 14, 55). In the present study, we decipher a mechanism of establishment and maintenance of PML NBs-associated quiescent HSV-1 genomes involving the apposition of the H3K9me3 mark on the viral chromatin by the HUSH-SETDB1-MORC2 repression complex. We show that his pathway is closely associated with, but independent of the formation of vDCP NBs. The HUSH-SETDB1-MORC2 complex is essential for the acquisition of the H3K9me3 chromatin mark on quiescent HSV-1 genomes and for their transcriptional repression. Questions may arise regarding the apparent absence or even contrary role of TASOR, at least in relation to the deposition of the H3K9me3 mark on quiescent HSV-1. These observations may need to be reconsidered in light of recent findings regarding the existence of a HUSH2 complex containing TASOR2, a paralog of TASOR, that may impact on the activity of HUSH (48). A functional HUSH-SETDB1-MORC2 complex is also essential for maintaining PML NBs-associated HSV-1 quiescence in neuronal cells, as suggested by the importance of MORC2 in the control of the stability of HSV-1 quiescence in hiPSDN. Given the essential role of MORC2 for the HUSH complex activity (35, 56), this suggests that the HUSH-SETDB1-MORC2 repression complex acts as a major epigenetic regulatory entity to maintain PML NBs-associated HSV-1 quiescence in neurons.

Accordingly, we found that MORC2 is a specific target of the viral protein ICP0 for degradation mediated by the proteasome. We and others have already published that ICP0 has multiple cellular targets to prevent the antiviral response (for reviews (57, 58)). Our results suggest that rather than targeting for degradation one of the three components of the HUSH complex, which would leave MORC2 to accomplish its activity independently of the HUSH complex, ICP0 induces the degradation of MORC2, which would affect the entire SETDB1-HUSH-MORC2 repressor complex. It is already known that ICP0 impacts on the stability of MORC3 a gatekeeper of the antiviral response (50, 59). The additional degradation of MORC2 mediated by ICP0 is thus of most importance and suggests that the virus has evolved to counteract the activity of the proteins of the entire MORC family, which comprises 4 members. This anticipates that proteins of the MORC family are major restriction factors that evolved to counteract HSV infection. Another important aspect regarding MORC2 is that it has been described together with MPP8 as a major component of neurons playing a major role in neurons homeostasis by repressing the protocadherin gene cluster, which play crucial roles in the development and function of the nervous system, particularly in neuronal connectivity and synaptic specificity (60). It is thus very interesting to see that a neurotropic virus such as HSV-1, which establishes latency in sensory neurons, has evolved as such as its viral protein essential for full reactivation, ICP0, targets MORC2 for degradation. SETDB1 is also known for its major involvement in neurons survival and brain development (61). The involvement of the HUSH-SETDB1-MORC2 entity in the acute control, then long-term maintenance, of one of the latency-associated molecular patterns of a neurotropic virus is thus of particular interest.

The association of the HUSH-SETDB1-MORC2 entity with vDCP NBs most likely links its epigenetic activity with the stability of the vDCP NBs. The HSV-1 ICP0 protein destabilizes PML NBs (62) and vDCP NBs (8), and has been shown to erase the H3K9me3 mark on viral chromatin upon HSV-1 lytic cycle and establishment/maintenance of latency (63, 64). We show that ICP0 is able to induce HSV-1 reactivation in quiescently infected iPSDN. This suggests a direct and/or indirect impact of ICP0 on the HUSH-SETDB1-MORC2 repression complex activity by destabilizing vDCP NBs.

The HUSH-SETDB1-MORC2 entity acting as a novel nuclear restriction pathway in concert with the activity of PML NBs, extends at the epigenetic level the intrinsic response of the cell for the control of the infection by a neurotropic virus (**Fig. 6e**). This pathway is a direct target of the activity of the viral protein ICP0, likely through the proteasomal degradation of MORC2 in addition to the destabilization of vDCP NBs, to promote reactivation of HSV-1. Stabilizing this important pathway would enable the control of neuronal damage caused by the spread of the virus in the nervous system following reactivation. Alternatively, destabilizing the HUSH-SETDB1-MORC2 entity might be considered as a complementary approach to proposed therapies designed to modify viral genomes to prevent their reactivation. As an example, viral genomes editing using meganucleases or CRISPR/Cas9 technologies has been proposed to eliminate latent HSV reservoirs (65–69). Provoking sub-reactivation of HSV-1 by inactivating the HUSH-SETDB1-MORC2 entity could then potentiate the use of these technologies, provided that it does not induce global epigenetic modifications that would put a threat on cellular defense mechanisms. On a different aspect, targeting HUSH for degradation has been discussed as part of a “shock and kill” strategy in HIV cure designed to eliminate reservoir cells responsible for rebounds when antiretroviral treatments are stopped (70). Our study establishes HUSH as a crucial factor in maintaining PML NBs-associated HSV-1 quiescence. Targeting HUSH degradation, moreover in immunocompromised patients, without controlling herpesvirus lytic cycle, e.g. by administration of anti-herpesviral drugs, could therefore compromise the integrity of patients. Indeed, this could lead to uncontrolled HSV-1 spread in the PNS and CNS potentially responsible for neuronal and/or ocular pathologies as a result of herpesvirus reactivation. In that context, the “block and lock” strategy to create a “deep latency” of HIV seems indeed more appropriate considering the likely broad involvement of the HUSH-SETDB1-MORC2 entity in the control of multiple latent viruses. Finally, the importance of the HUSH-SETDB1-MORC2 repression complex together with PML NBs to control a genuine unintegrated virus, such as HSV-1, could be broaden to other viruses remaining latent/quiescent as episomes in the nucleus of infected cells.

## Methods

### Virus strains

The HSV-1 mutant *in*1374 is derived from the 17 *syn* + strain and expresses a temperature-sensitive variant of the major viral transcriptional activator ICP4 (71). *In*1374 is derived from *in*1312, a virus derived from the VP16 insertion mutant *in*1814 (72), which additionally carries a deletion/frameshift mutation in the ICP0 open reading frame (73) and contains an HCMV-*lacZ* reporter cassette inserted into the UL43 gene of *in*1312 (74). This virus has been used and described previously (4, 6). *In*1374 was grown and titrated at 32°C in the presence of 3 mM HMBA (75). For simplicity, *in*1374 will be renamed lqHSV-1 (for latent/quiescent HSV-1).

### Cells

Cells were tested regularly for mycoplasma contaminations; experiments were only performed on non-contaminated cells. Human BJ primary foreskin fibroblasts (ATCC, CRL-2522), noted hFC hereafter and in the main text, human embryonic kidney (HEK 293T, ATCC CRL-3216, kind gift from M. Stucki, University Hospital Zürich) cells, human bone osteosarcoma (U2OS) cells (kind gift from R. Everett), and baby hamster kidney (BHK-21) cells (kind gift from R. Everett) cells were grown in Dulbecco’s Modified Eagle’s Medium (DMEM) (Sigma-Aldrich, D6429) supplemented with 10% fetal bovine serum (Sigma, F7524), L-glutamine (1% v/v), 10 U/mL penicillin, and 100 mg/mL streptomycin (Sigma-Aldrich, P4458). hFC division is stopped by contact inhibition. Therefore, to limit their division, cells were seeded at confluence before being infected at a multiplicity of infection (m.o.i.) of 3, and then maintained in 2% serum throughout the experiment.

Human induced pluripotent stem (hiPS) cells (clone AG08C5) were derived at the Marseille Medical Genetics (MMG) (76), were maintained in culture on plates coated with Matrigel (Corning, 354277) in mTeSR+ (Stemcell Technologies, 05825). The medium was changed every 2 days. Cells were passaged using tryple (Gibco, 354277), washed, and replated at 2.10e5 cells. Revitacell (Rock inhibitor) was added during the passage in order to promote cell survival.

### Generation of sensory neurons

Differentiation of hiPSC into sensory neurons was performed according to (77). Briefly, hiPSC colonies at 60% confluence were dissociated using tryple and pelleted at 800 rpm for 4 minutes (room temperature). To generate sensory neurons, the cells were maintained on Matrigel-coated plates in DMEM/F12 with defined supplements. Cultures were supplemented from day 1 to day 10 with 10% KSR, 1% penicillin/streptomycin, 0.3 mM LDN-193189, 2 mM A83-01, 6 mM CHIR99021, 2 mM RO4929097, and 3 mM SU5402. Retinoic acid 0.3 mM was included in all cultures. The medium was refreshed every 2 days. The mitotic inhibitors fluorodeoxyuridine was added at day 6 until day 10. The medium was then replaced with fresh medium without fluorodeoxyuridine. Once day 10 is reached, cultures were maintained in neurobasal medium supplemented with 10 ng/ml neurotrophin 3 (NT3), 20 ng/ml NGF with half of the medium refreshed every other day for up to 2 weeks.

### Nociceptive properties of hiPSDN

Sensory neurons with nociceptive activity are characterized by their ability to release calcitonin gene-related peptide (CGRP) following capsaicin treatment. Neurons were treated overnight with either 1, 10, 100uM of Capsaicin or KCl. Supernatants were then speedvac and analyzed by « alpha-CGRP human enzyme immunoassay » (Cat. No. S-1199 from Peninsula Laboratories,San Carlos, CA, USA). All was performed according to the manufacturer instructions. Analysis of the results was performed with Prism software, by applying the “log (inhibitor) vs. response -- Variable slope (four parameters)” model.

### Reagents

Drugs and molecules used for cell treatments are described below (duration is mentioned in the main text)

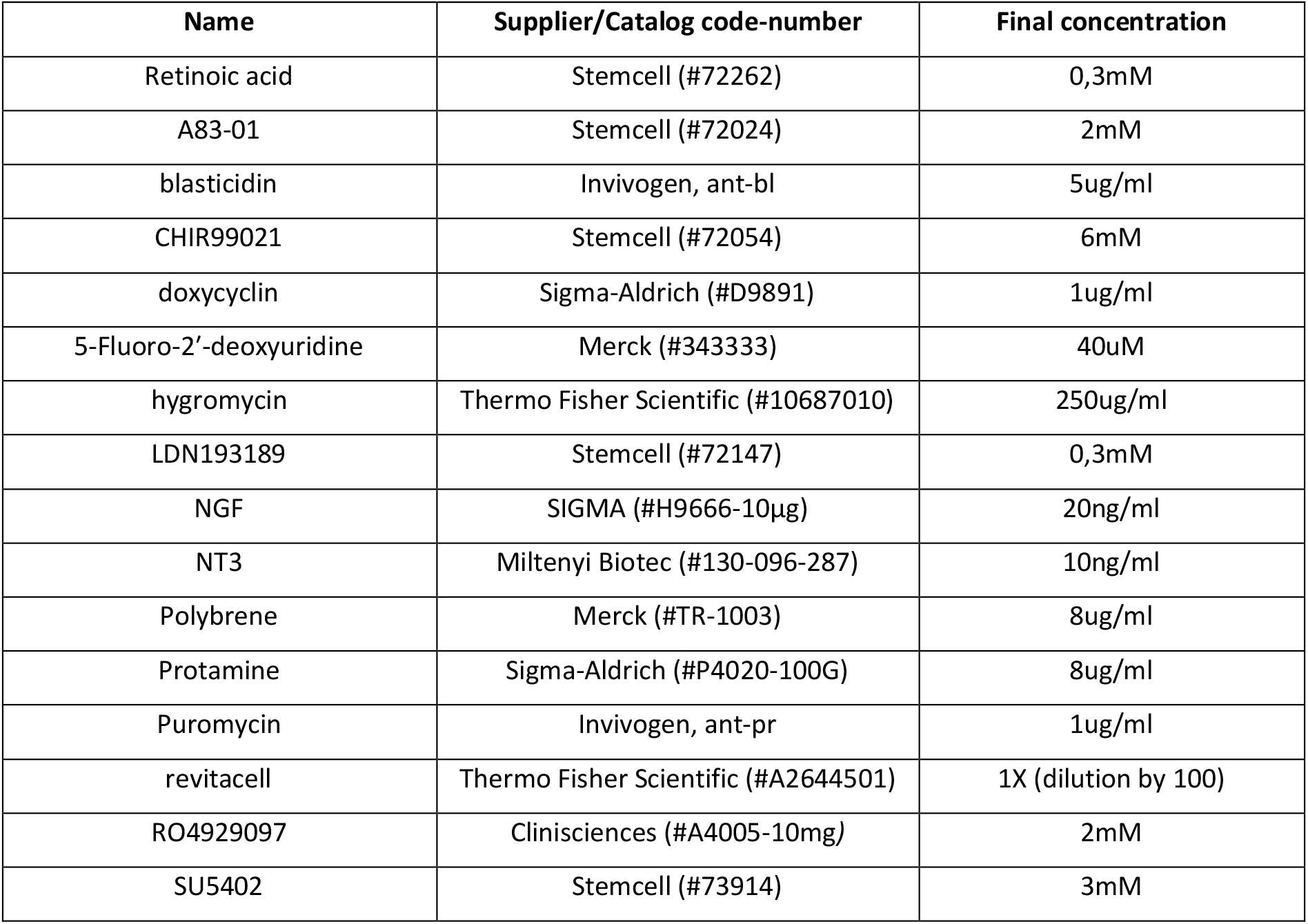

### Electrophysiology

Whole-cell patch-clamp recordings were performed at 30 °C using a Multiclamp 700B amplifier. Electrodes had a tip resistance of 5–7 Mohm when filled with intracellular solution (140mM CsCl, 1mM MgCl2, 1mM CaCl2, 10mM EGTA, 1mM BAPTA, 10mM HEPES and 4mM MgATP). The perfusion medium contained 145mM NaCl, 2.4mM KCl, 10mM HEPES, 10mM D-glucose, 2mM CaCl2 and 2mM MgCl2, and was supplemented when necessary with 1 µM TTX, to test the presence of voltage gated sodium channels. Recordings were filtered at 2– 5 kHz and acquired at 5–10 kHz. Currents were recorded in voltage clamp mode at a holding potential of −80 mV. 250 ms and 10 mV, incremental steps were preformed to test the current-voltage relationship and elicit voltage gated sodium channels opening. Cells in which the access resistance changed by 20% during the recording or exceeded 20 Mohm were discarded. Data were acquired and analyzed with clampex 10 software (Axon).

### Lentivirus production and establishment of cell lines

hFC or hiPSDN cells stably expressing transgenes or shRNA were obtained by lentiviral transduction. Briefly, plasmids of interest (see plasmids section) were co-transfected with psPAX.2 (Addgene #12260) and pMD2.G (Addgene #12259) plasmids with a ratio of 3:2:1 by the calcium phosphate method into HEK 293T cells to package lentiviral particles. After 48 h, supernatant containing replication-incompetent lentiviruses was filtered and applied for 24 h on the target cells in a medium containing polybrene 8 μg/mL (Sigma) (78) in the case of hFC or protamine 8ug/ml (Sigma) in the case of hiPSDN. Stable transfectants were selected with Blasticidin S (5 μg/mL, Invivogen), puromycin (1 μg/mL, Invivogen), or hygromycin (250 ug/mL, thermo), and a polyclonal population of cells was used for all experiments

### Plasmids

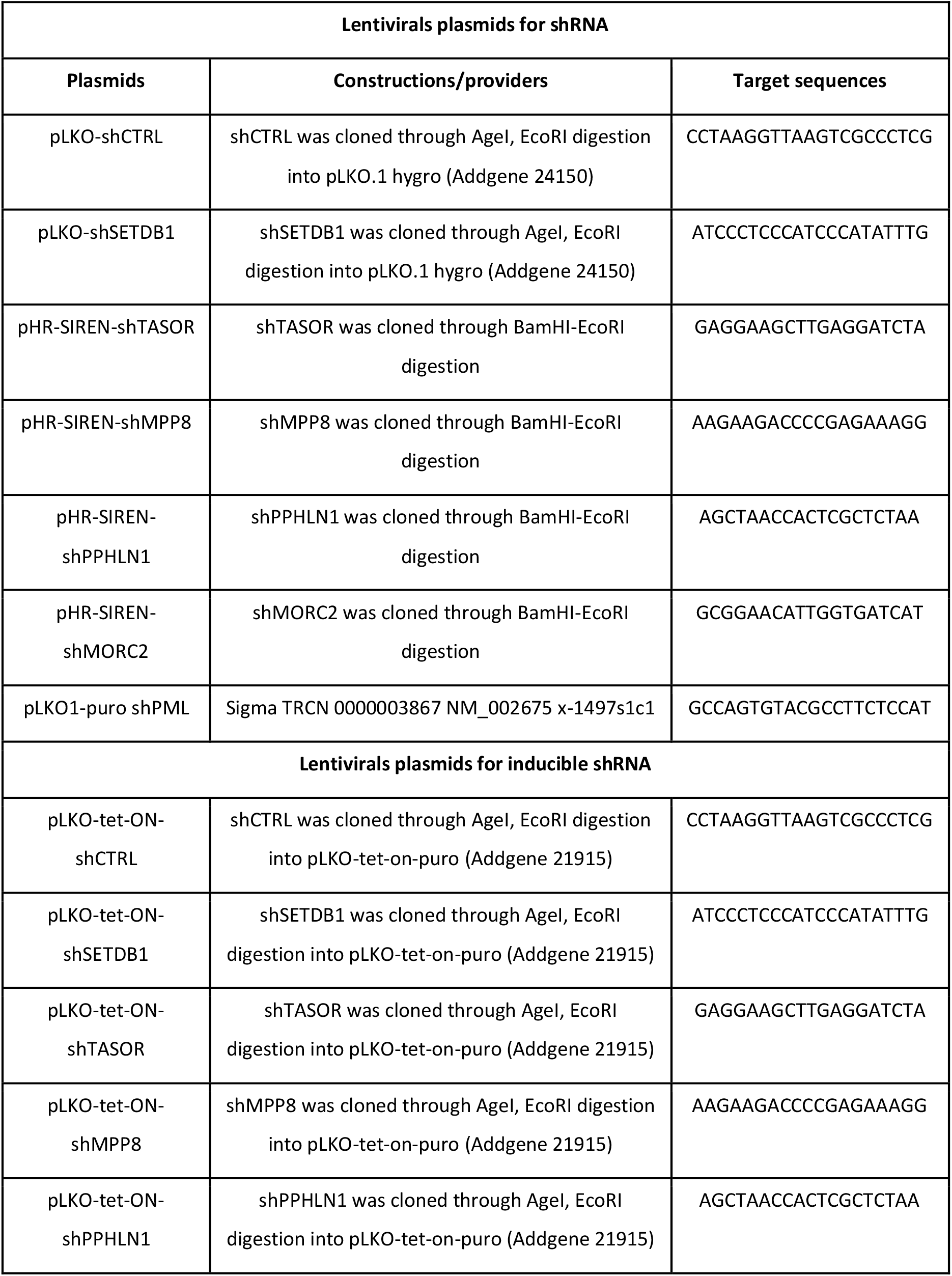
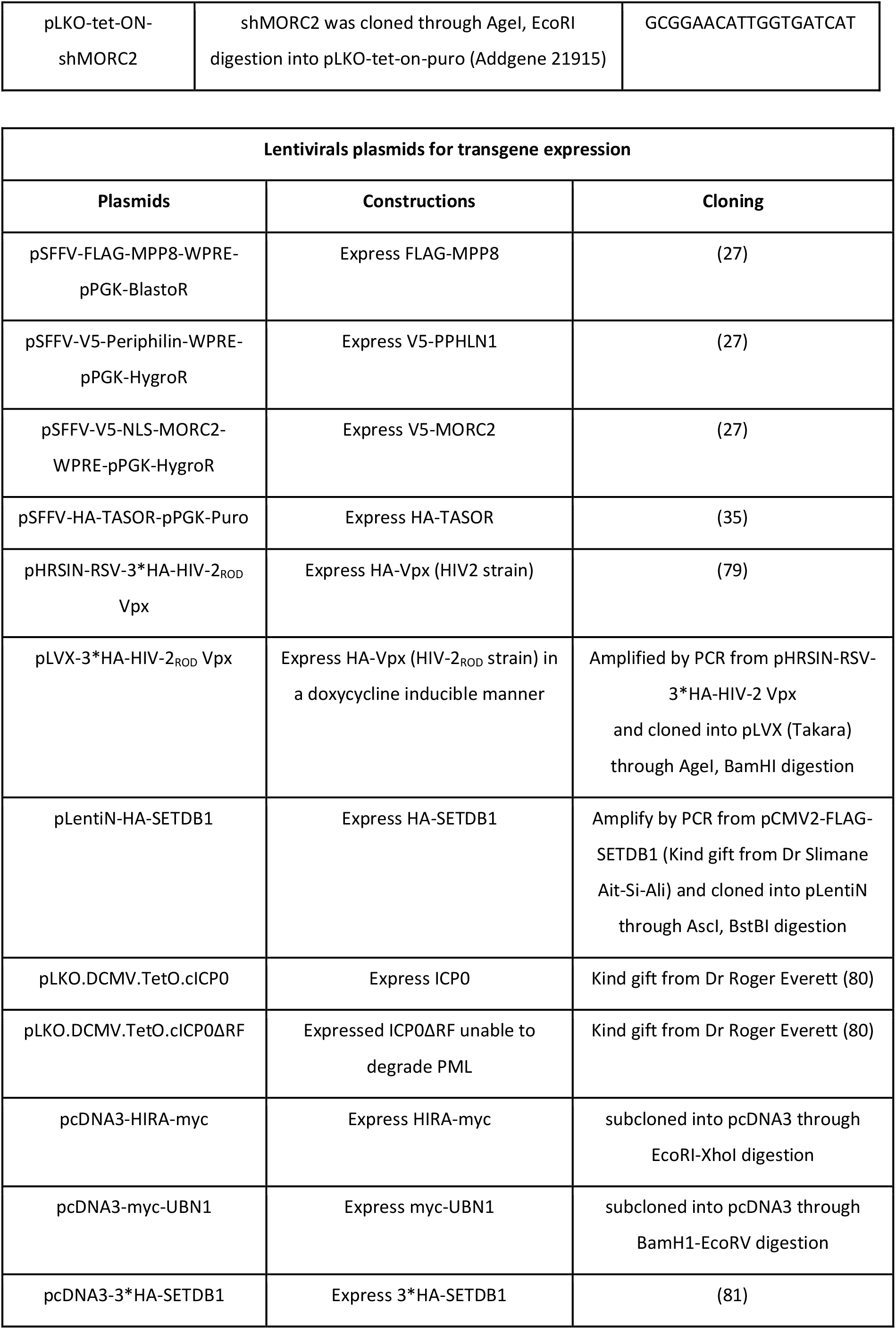

### Immunofluorescence

Cells were seeded on either 24-well plate containing one coverslip or directly into 8-wells labteck. Cells were fixed for 10 min with 4% paraformaldehyde, washed three times with PBS, then permeabilized for 5 min in a PBS solution containing 0.5% Triton X-100. Primary antibodies were diluted in PBS containing 3% newborn goat serum (see antibodies section). After incubation at room temperature for 1 h, coverslips were washed at least three times with PBS plus 3% newborn goat serum, and then incubated with goat anti-mouse or anti-rabbit (H+L) cross-absorbed secondary antibodies Alexa-488, Alexa-555 or Alexa-647 (Invitrogen). After a further 30-min incubation, coverslips were washed three times with PBS plus 3% newborn goat serum. Following immunostaining, nuclei were stained with DAPI (Invitrogen Life Technologies, D1306) diluted in PBS at 0,1μg/mL for 5 minutes at RT°C. Coverslips were mounted in Fluoromount-G (SouthernBiotech, 0100-01) and stored at 4°C before observation.

### DNA-FISH and immuno-FISH

HSV-1 DNA FISH probes consisting of cosmids 14, 28 and 56 (82) comprising a total of ∼90 kb of the HSV-1 genome were labeled by nick-translation (Invitrogen) with dCTP-Cy3 (GE Healthcare) and stored in 100% formamide (Sigma). The DNA-FISH and immuno-DNA FISH procedures have been described previously (3, 83). Briefly, infected cells were fixed in 2% PFA. Infected cells were then permeabilized in 0.5% Triton X-100. Heat-based unmasking was performed in 100 mM citrate buffer, and sections were post-fixed using a standard methanol/acetic acid procedure and dried for 10 min at RT. DNA denaturation of the section and probe was performed for 5 min at 80°C, and hybridization was carried out overnight at 37°C. Sections were washed 3 x 10 min in 2 x SSC and for 3 x 10 min in 0.2 x SSC at 37°C, and nuclei were stained with DAPI (Invitrogen, Life Technologies, D1306). All sections were mounted under coverslips using Fluoromount-G (SouthernBiotech, 0100-01)

For immuno-DNA FISH (FISH-IF), cells were treated as described for DNA-FISH up to SSC washing. Cells were then treated with PBS-Normal Goat Serum to saturate non-specific site for 1h at RT. Cells were then incubated with the primary antibody 1h (see antibodies section). After three washes, secondary antibody was applied for 1 h. Following immunostaining, nuclei were stained with DAPI (Invitrogen Life Technologies, D1306) diluted in PBS at 0,1μg/mL for 5 minutes at RT°C. Coverslips were mounted in Fluoromount-G (SouthernBiotech, 0100-01) and stored at 4°C before observation.

### Microscopy, imaging, and quantification

Images were acquired with the Axio Observer Z1 inverted wide-field epifluorescence microscope (100X or 63X objectives/N.A. 1.46 or 1.4) (Zeiss) and a CoolSnap HQ2 camera from Photometrics. Identical settings and contrast were applied for all images of the same experiment to allow data comparison through ImageJ.

### ChIP and quantitative PCR

Cells were fixed with methanol-free formaldehyde (#28908, Thermo Fisher Scientific) 1% for 5 min at RT, and then glycine 125 mM was added to arrest fixation for 5 min. After two washes with ice-cold PBS, the cells were scraped and resuspended in “Lysis Buffer” (10% glycerol, 50mM HEPES pH7,5; 140mM NaCl; 0,8% NP40;0,25% Triton; 1mM EDTA, Protease Inhibitor Cocktail 1X (PIC) (Complete EDTA-free; Roche) and incubated for 10 min at 4°C under shaking. The cells were subsequently washed in “Wash buffer” (200mM NaCl; 20mM Tris pH8; 0,5mM EGTA; 1mM EDTA, PIC 1X) for 10 min at 4°C under shaking then were resuspended and centrifuged twice during 5 min 1700g at 4°C in “Shearing Buffer” (10mM Tris pH7,6; 1mM EDTA; 0,1%SDS; PIC 1X). Finally, nuclei were resuspended in 1mL of “Shearing Buffer” and were sonicated with a S220 Focused-ultrasonicator (Covaris) (Power 140W; Duty Off 10%; Burst Cycle 200). Eighty-five μL of the sonication product were kept for the input, 50 μL for analyzes of the sonication efficiency, and 850 μL diluted twice in IP buffer 2X (300mM NaCl, 10mM Tris pH8; 1mM EDTA; 0,1% SDS; 2% Triton) for ChIP. Either two or five ug of Ab were added and incubated overnight at 4°C (see antibodies section). The next day, 50 µL of agarose beads coupled to protein A (Millipore 16–157) or G (Millipore 16–201) were added for 2 h at 4°C under constant shaking. Alternatively, agarose beads were coated with either 2 or 5 ug of antibody before the addition of the sonication product and incubated overnight at 4°C. For V5-tagged proteins, anti-V5 agarose affinity gel (Sigma-Aldrich, A7345) were incubated overnight at 4°C. Beads were then successively washed for 5 min at 4°C under constant shaking once in “low salt” (0.1% SDS, 1% Triton X-100, 2 mM EDTA, 20 mM Tris HCl pH 8.0, 150 mM NaCl) buffer, once in “high salt” (0.1% SDS, 1% Triton X-100, 2 mM EDTA, 20 mM Tris HCl pH 8.0, 500 mM NaCl) buffer, once in “LiCl” (0.25 mM LiCL, 1% NP40, 1% NaDOC, 1 mM EDTA, 10 mM Tris HCl pH 8.0) buffer, and twice in TE (10 mM Tris pH 8.0, 1 mM EDTA) buffer. Chromatin-antibody complexes are then eluted at 65°C for 30 min under constant shaking with 200 μL of elution buffer (1% SDS, 0.1 M NaHCO3). Input and IP products were de-crosslinked overnight at 65°C with 20 mg/mL of RNAse (Sigma) then treated for 2 hours at 55°C with 20 mg/mL of proteinase K (Sigma). DNA was then purified by phenol-chloroform/ethanol precipitation, resuspended in water, and kept at −20°C until use for qPCR. Quantitative PCRs were performed using the KAPA SYBR qPCR Master Mix (SYBR Green I dye chemistry) (KAPA BIOSYSTEMS, KK4618) and the CFX connect apparatus (Bio-Rad). Primers used are listed below:

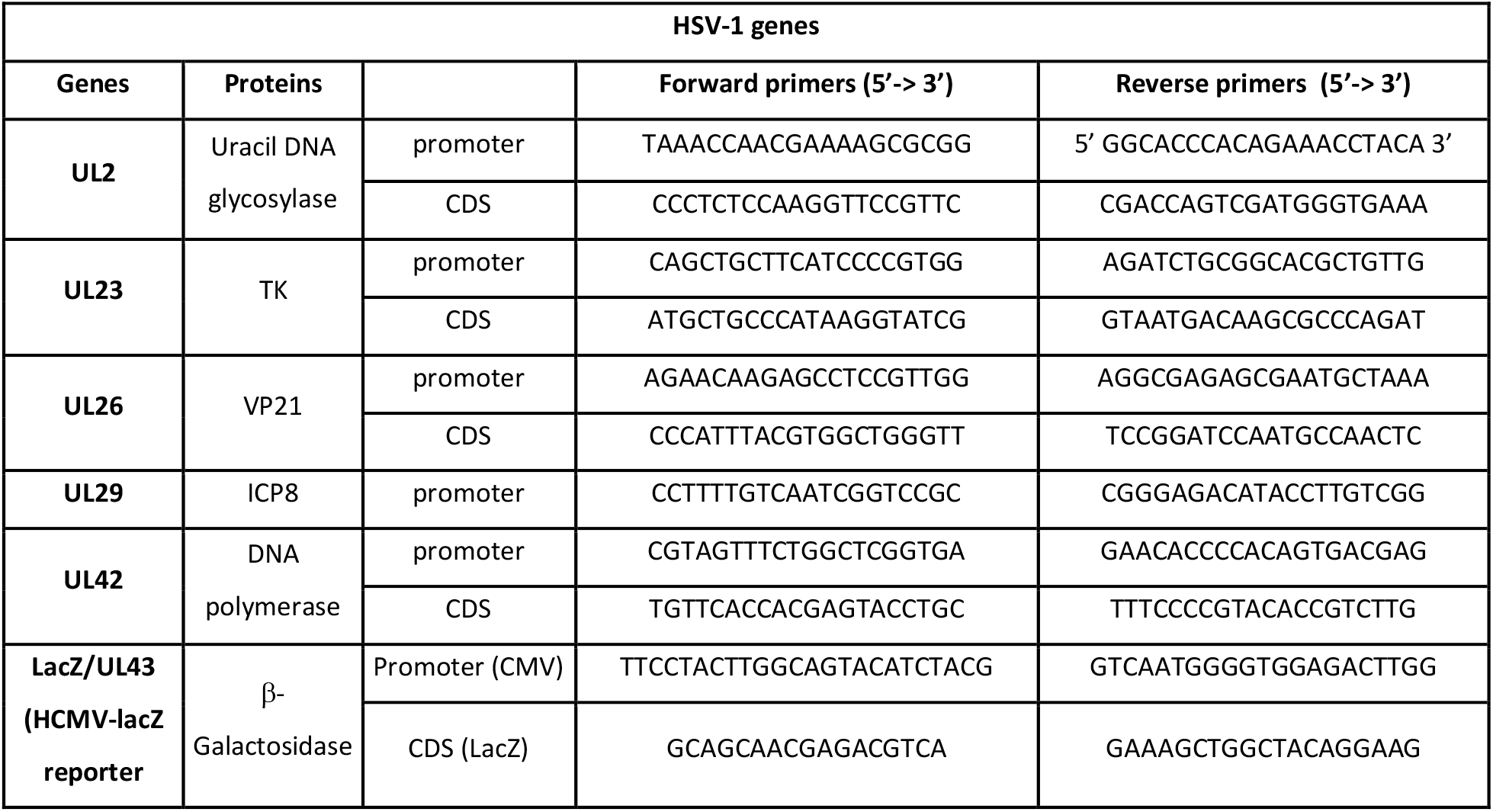
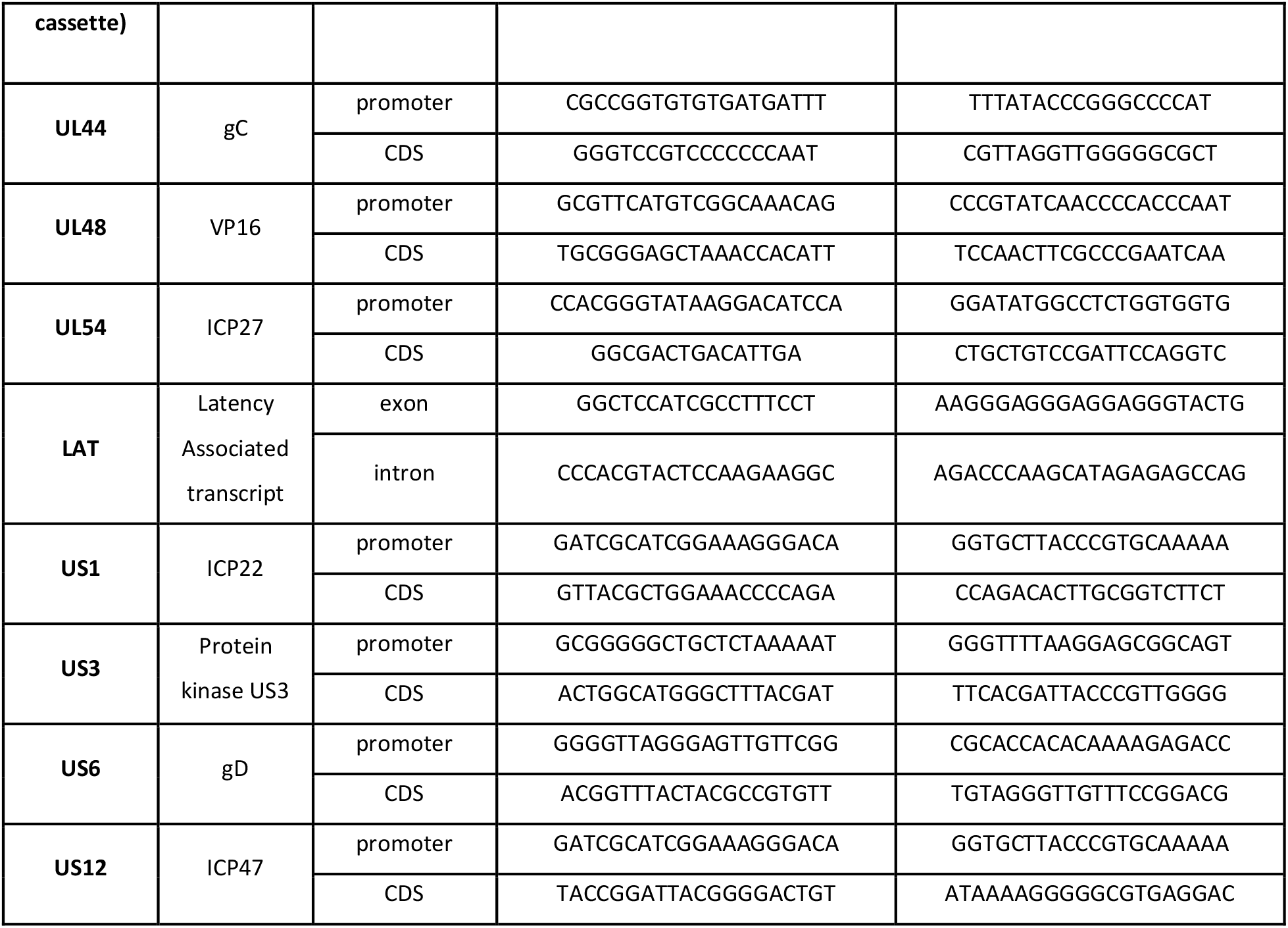

#### ChIP-Seq analysis

hFC were transduced with fresh lentiviruses encoding shRNAs against the HUSH complex, MORC2 or SETDB1 (see plasmids section) for 48h to allow the depletion of the specific proteins, before being infected with HSV-1 *in1374* at an MOI of 3 for 24h. Cells were then crosslinked and processed for ChIP as described previously (Kleijwegt et al., 2023). We used 20 μL of protein A magnetic dynabeads (Invitrogen, 10001D) for immunoprecipitation with 2 μg of the H3K9me3 antibody (see antibodies section). After ChIP, librairies were prepared with 10ng of DNA (input/ChIP) using the NEBNext® UltraTM II DNA Library Prep Kit for Illumina. DNA librairies were then sequenced using the paired-end method (2x150pb) on the Element Biosciences Aviti (HELIXIO, Clermont-Ferrand, France). Raw reads were filtered by trimming Illumina adapters and removing low-quality reads using Trimmomatic (v.0.39) with default parameters. Trimmed reads (containing more than 36 nucleotides) were aligned to the human (hg38) reference genome concatenated with the HSV-1 genome (GCF_000859985.2.fa) using bowtie2 (v2.5.1). Sequence alignment map files, including uniquely mapped and multi-mapped reads, were transformed and sorted into BAM files using SAMtools (v.1.15.1). Duplicate reads were removed using the MarkDuplicates function of Picard (v.2.23.5) (https://github.com/broadinstitute/picard) with REMOVE_DUPLICATES = true. Alignment files in the BAM format were converted to read coverage files in Bigwig format using deepTools bamCoverage (v.3.5.0) with RPGC normalization and 10 bp window size. Bigwig files comparing ChIP enrichment as a log2 fold change over input or over shCtl_NI were generated using deepTools bigwigCompare (v.3.5.0). Normalised bigwig files were displayed in IGV (v.2.15.1). Peak calling was performed using macs2 (v.2.2.7.1) in a broad mode with a broad cutoff of 0.01. After peak calling, peaks were separated between human and HSV-1 genomes to allow analysis on each genome separately. Peak annotation was performed in Rstudio (v.2023.12.0+369) with the readPeakFile, annotatePeak and plotAnnoBar functions of the ChIPseeker package. Heatmaps and profile plots were generated with deepTools computeMatrix, plotHeatmap and plotProfile commands (v.3.5.0). We used ChIP-Seq data against H3K9me3 (Table S1 from Becker et al., 2017, GSE16368) to plot H3K9me3 density across large heterochromatin regions previously identified in hFC. The LINEs sequences in a .bed format were downloaded from the RepeatMasker UCSC genome browser (GRCh38/hg38). Subtypes of LINEs were filtered in Bash using the grep command to allow creation of heatmaps on specific subtypes of LINEs (eg L1PA6). Matrices generated with computeMatrix on LINEs .bed files were imported in R in order to calculate the mean signal coverage per LINE subtype. Results were plotted with the ggplot function of R. After verifying that the data does not follow a normal distribution with a Kolmogorov-Smirnov test, we tested whether the mean signal coverage of two samples differ significantly from each other by using the non-parametric Wilcoxon rank-sum test. The FeatureCounts command of the subread package (v.2.0.1) was used to count reads on the HSV-1 genes using the GCF_000859985.2_HSV-1.gtf file. All bioinformatic analyses were performed on the Institut Français de Bioinformatique (IFB) core cluster through JupyterHub. Figures were made with RStudio (v.2023.12.0+369) or with Inkscape (v.1.0.2 (e86c8708, 2021-01-15)).

### siRNA transfections

Transfections of hFC with siRNAs was performed using Lipofectamine RNAiMAX and following the supplier’s procedure (Thermo Fisher Scientific). The following siRNAs were used at a final concentration of 40 nM for 48 h: siCTRL (siluc) 5’-CGUACGCGGAAUACUUCGA (84), siSETDB1 : 5’ACCCGAGGCUUUGCUCUUA (27), siPML: 5’AGATGCAGCTGTATCCAAG (85).

### Reactivation procedures

#### hFC

Cells were transduced with pLKO-tet-On-shRNA to produce hFC-ishRNA stably and inducibly expressing the ishRNA (selection puro 1ug/mL). FBS Tetracycline free (Clinisciences, FBS-TET-12A) was used to prevent ishRNA expression. hFC-ishRNA were infected with lqHSV-1 for 4 days at 38.5 °C to stabilize the formation of vDCP NBs. Then, the expression of shRNA was induced by the addition of doxycycline (1ug/mL) in the medium. Cells were incubated at 32 °C the permissive temperature for *lq*HSV-1 (see section virus). From day 8 to day 10 after addition of doxycycline, the cells were fixed to proceed to immuno-FISH analyses or RNA were extracted to perform RT-qPCR (see section RT-qPCR).

#### HiPSDN

HiPSDN were cultured for at least 24 hours without antimitotic agents prior to infection with lqHSV-1. HiPSDN were infected with the indicated virus at 3.10e5 PFU/wells for 1h30 at 37 °C. Post-infection, inoculum was replaced with neurobasal containing NGF (20ng/ml), NT3 (10ng/ml). After 4 days at 38°C, neurons were transduced with the indicated lentivirus (see lentivirus section). The next day, transduced neurons were put at 32°C (the permissive temperature for replication) during the indicated times. Human Serum (1%) was added to the media to limit cell-to-cell spread. Reactivation was quantified by RT-qPCR of HSV-1 lytic mRNAs isolated from the cells in culture. Same experiment was performed but without Human serum to recover the supernatant. Reactivation was quantified by the production of infectious viral particles through supernatant titration.

### RT-qPCR

RNAs were extracted according to The Qiagen RNeasy Mini Kit manufactured protocol (Qiagen, Valencia, CA). RNAs were treated 30mn at 37°C with DNAseI (thermo fisher). Then 1ug was taken to perform RT with RevertAID Reverse Transcriptase (thermo fisher). cDNA was then resuspended in water and kept at −20°C until use for qPCR. Quantitative PCRs were performed using the KAPA SYBR qPCR Master Mix (SYBR Green I dye chemistry) (KAPA BIOSYSTEMS, KK4618) and the CFX connect apparatus (Bio-Rad). Primers used are listed below:

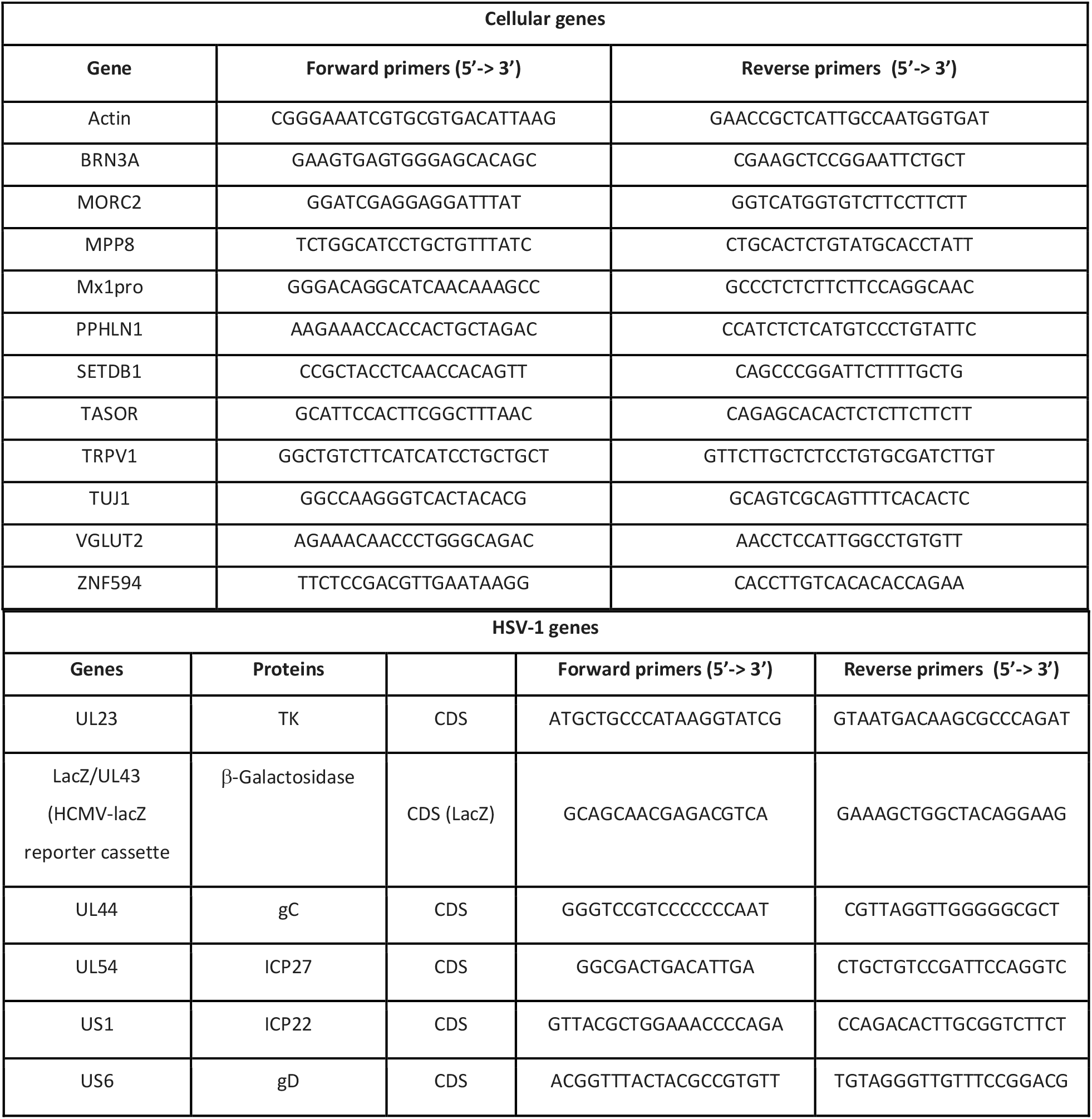

### Western blotting

RIPA extracts were obtained by lysing the cells in RIPA buffer (50mM Tris-HCl pH 7.5, 150mM NaCl, 0,5% Na-Deoxycholate, 1% NP-40, 0,1% SDS, 5mM EDTA) supplemented with 1X protease inhibitor cocktail (PIC) for 20min on ice. After incubation, RIPA extracts were centrifugated for 10min at 16000g at 4°C and supernatants were recovered and diluted with 4X LSB (250mM Tris-HCl pH 6.8, 40% glycerol, 8% SDS, bromophenol blue). Protein extracts were boiled for 10 minutes and then loaded on polyacrylamide gels (TGX Stain-Free FastCast gels (Bio-Rad, 1610181/1610183) or home-made gels) for electrophoresis. Stain-free technology or Ponceau 0,1% (Sigma-Aldrich, P7170) were used to reveal proteins transferred on nitrocellulose membrane with the Trans-Blot Turbo Transfer System (Bio-Rad, 1704150). Membranes were blocked in PBS 0.1% Tween20 (PBST) 5% milk for 30 minutes at RT°C and incubated with primary antibodies diluted in PBST or PBST 5% BSA O/N at 4°C (see antibodies section). After washing the membranes twice in PBST, they were incubated in secondary antibodies conjugated with horseradish peroxidase (HRP) (Dako, P0399, P0447 and P0450) diluted according to supplier recommendations in PBST for 1 hour at RT°C. Signal was revealed on ChemiDoc Imaging System (Bio-Rad) by using Amersham ECL Prime Western Blotting Detection Reagent (GE Healthcare Life Sciences, RPN2236) or Clarity Max Western ECL Blotting Substrate (Bio-Rad, 1705062).

### Antibodies

The following primary antibodies were used:

#### For ChIP/ChIP-seq*

| Proteins | Supplier | Catalog number | Species |
| --- | --- | --- | --- |
| Flag | Sigma | f1804 | Mouse monoclonal |
| HA | Abcam | ab9110 | Rabbit polyclonal |
| H3 | Abcam | 1791 | Rabbit polyclonal |
| H3K9me3* | Abcam | ab8898 | Rabbit polyclonal |
| Immunoglobulin G | Diagenode | Kch-504-250 | Rabbit polyclonal |
| MPP8 | Proteintech | 16796-1-AP | Rabbit polyclonal |
| V5 | Sigma | A7345 | Mouse monoclonal |

#### For IF and/or FISH-IF

For IF and/or FISH-IF:
| Proteins | Supplier | Catalog number | Species | Dilution |
| --- | --- | --- | --- | --- |
| Flag | Sigma | f1804 | Mouse monoclonal (clone M2) | 1/200 |
| HA | Abcam | ab9110 | Rabbit polyclonal | 1/1000 |
| ICP4 | Viruzys | P1101 | Mouse monoclonal (clone H943) | 1/250 |
| MORC2 | Sigma | HPA000436 | Rabbit polyclonal | 1/250 |
| MPP8 | Proteintech | 16796-1-AP | Rabbit polyclonal | 1/400 |
| NeuN | Millipore | MAB377 | Mouse monoclonal (clone A60) | 1/100 |
| NF200 | Sigma | N4142-.2ML | Rabbit polyclonal | 1/200 |
| PML | Santa cruz | Sc-966 | Mouse monoclonal (PG-M3) | 1/250 |
| PML | Sigma | PLA0172 | Rabbit polyclonal | 1/5000 |
| PPHLN1 | Sigma | HPA038902 | Rabbit polyclonal | 1/150 |
| SETDB1 | Abcam | ab107225 | Mouse monoclonal | 1/250 |
| SETDB1 | Proteintech | 11231-1-AP | Rabbit polyclonal | 1/250 |
| TASOR | Sigma | HPA006735 | Rabbit polyclonal | 1/200 |
| TUJ1 | BioLegend | PRB-435P-050 | Rabbit polyclonal | 1/1000 |
| TRPV1 | Alomone labs | ACC030 | Rabbit polyclonal | 1/200 |
| V5 | Thermofisher | R960-25 | Mouse monoclonal | 1/1000 |

All secondary antibodies were Alexa Fluor-conjugated and 1 were raised in goats (Invitrogen).

#### For WB

1 All secondary antibodies were Alexa Fluor-conjugated and were raised in goats (Invitrogen). 2 For WB:
| Proteins | Supplier | Catalog number | Species |
| --- | --- | --- | --- |
| Actin | Sigma | A2066 | Rabbit polyclonal |
| α-actinin | Millipore | 05384 | Mouse (clone AT6/172) |
| MORC2 | Bethyl laboratories | A300-149A | Rabbit polyclonal |
| MPP8 | Proteintech | 16796-1-AP | Rabbit polyclonal |
| PPHLN1 | Sigma | HPA038902 | Rabbit polyclonal |
| SETDB1 | Proteintech | 11231-1-AP | Rabbit polyclonal |
| TASOR | Sigma | HPA006735 | Rabbit polyclonal |
| α-Tubulin | Sigma | T6199 | Mouse monoclonal (clone DM1A) |
| H3 | Abcam | 1791 | Rabbit polyclonal |
| H3K9me3 | Abcam | ab8898 | Rabbit polyclonal |
| HA | Abcam | ab9110 | Rabbit polyclonal |

All secondary antibodies were HRP-conjugated and were raised in goats (Sigma).

## Acknowledgements

We thank Prof. Roger Everett (Center for Virus Research, University of Glasgow, UK) for the *in*1374/lqHSV-1 virus and ICP0-expressing lentivirus vectors, Dr. Slimane Ait-Si-Ali (Epigenetic and cell fate, University Paris Diderot, France) for SETDB1 constructs, Prof. Delphine Poncet (INMG-PGNM) for advices on statistical analyses, Dr. Thomas Simonet (INMG-PGNM) for precious advice on bioinformatic analyses.

INMG-PGNM laboratory is funded by grants from the Centre National de la Recherche Scientifique (CNRS), Institut National de la Santé et de la Recherche Médicale (Inserm), University Claude Bernard Lyon 1, and AFM-téléthon. This work was supported by the French National Agency for Research-ANR through grants ANR-18-CE15-0014-01, ANR-20-COV9-0004-01, ANR-21-CE17-0018, LabEX DEV2CAN (ANR-10-LABX-61), ligue contre le cancer, and ANRS (ECTZ188412, ECTZ159208), AFM-téléthon (MyoNeuRALP2). PL team is member of the Groupements de Recherche, ‘’Dynamique des interactions entre chromatines virale et cellulaire’’ (DYNAVIR, GDR2194), and ‘’Architecture et Dynamique du Noyau & des Génomes’’ (ADN&G) funded by the CNRS. PJL is supported by a Wellcome Trust Principal Research Fellowship ((210688/Z/18/Z). PL is a CNRS Research Director.

## Author contributions

S.R., T.E., F.J., A.C., P.J.L., P.L. conceived the study. S.R., T.E., F.J., A.C., M. L., F.M., P.J.L., P.L. designed experiments and interpreted data. S.R., T.E., F.J., A.C., R.N., O.B., C.S., M.G., C.C., P.T., N.O., O.H., O.P., Y.G., M.L. performed experiments. F.M. developed the hiPSC and contributed to the hiPSDN differentiation protocol. A.C., S.B. contributed to the development and production of lentiviruses. O.P. performed the electrophysiological experiments for hiPSDN. S.R., T.E., F.J., A.C., P.J.L. and P.L. wrote the paper. All authors reviewed and edited the manuscript.

## Data availability statement

The datasets generated and/or analysed during the current study are available from the corresponding author on reasonable request.

---

**Figure S1.**
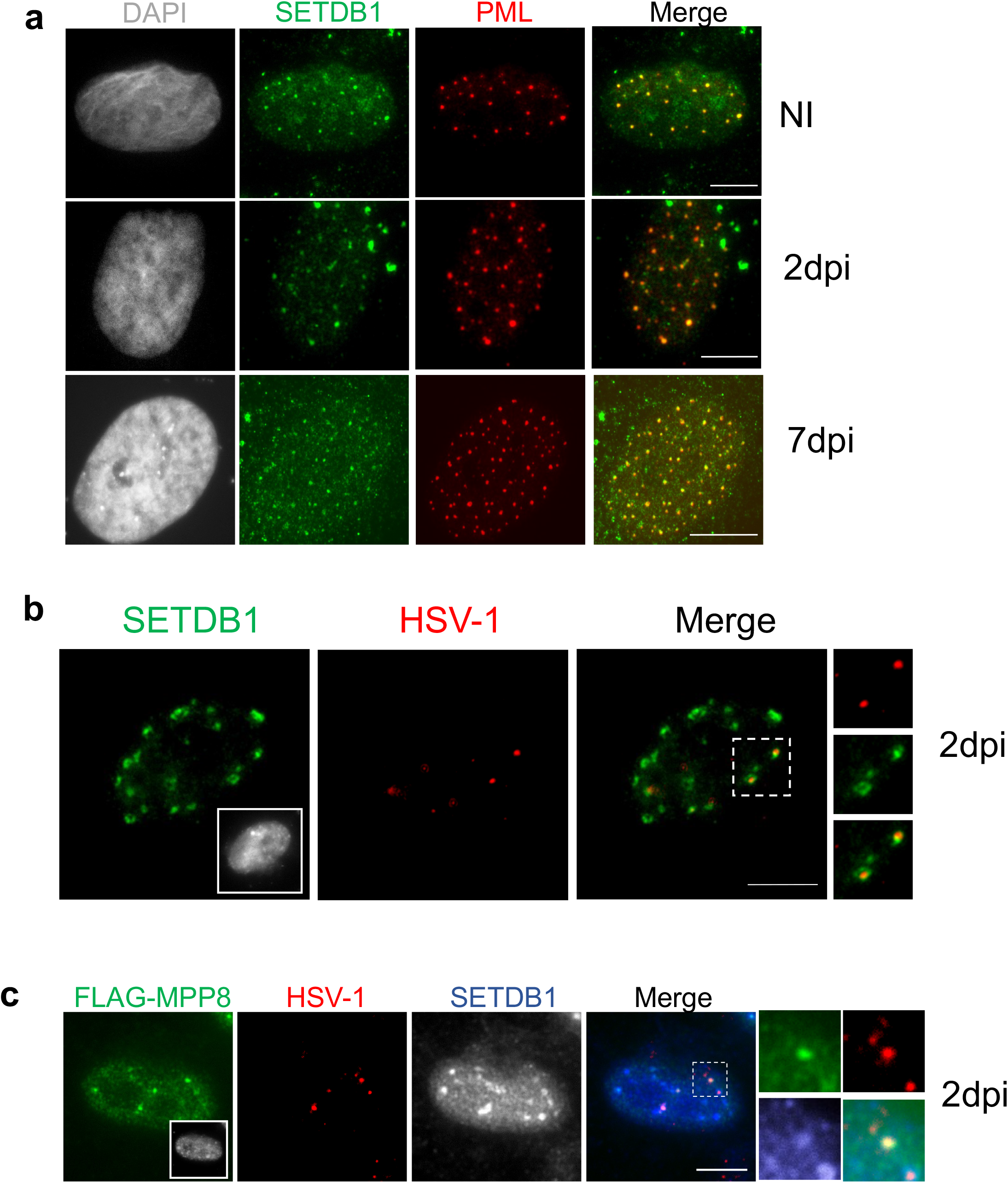

**Supplementary Fig.2.**
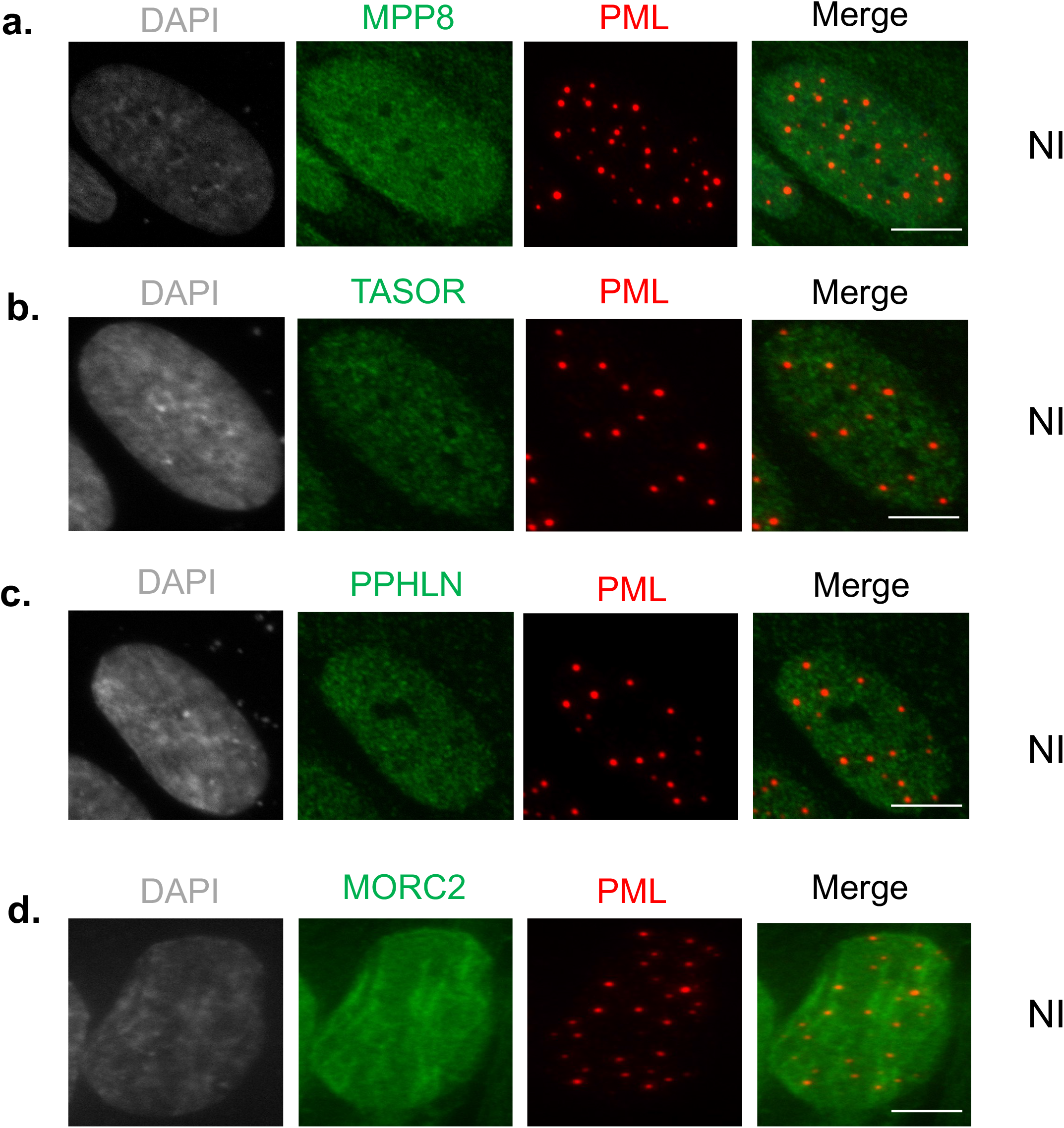
Endogenous staining of MPP8, TASOR, PPHLN1, MORC2 and PML NBs. **a to d,** Immunofluorescences on hFC of endogenous proteins of interest (green) co-stained with PML (red). Scale bars represent 5µm.

**Supplementary Fig.3.**
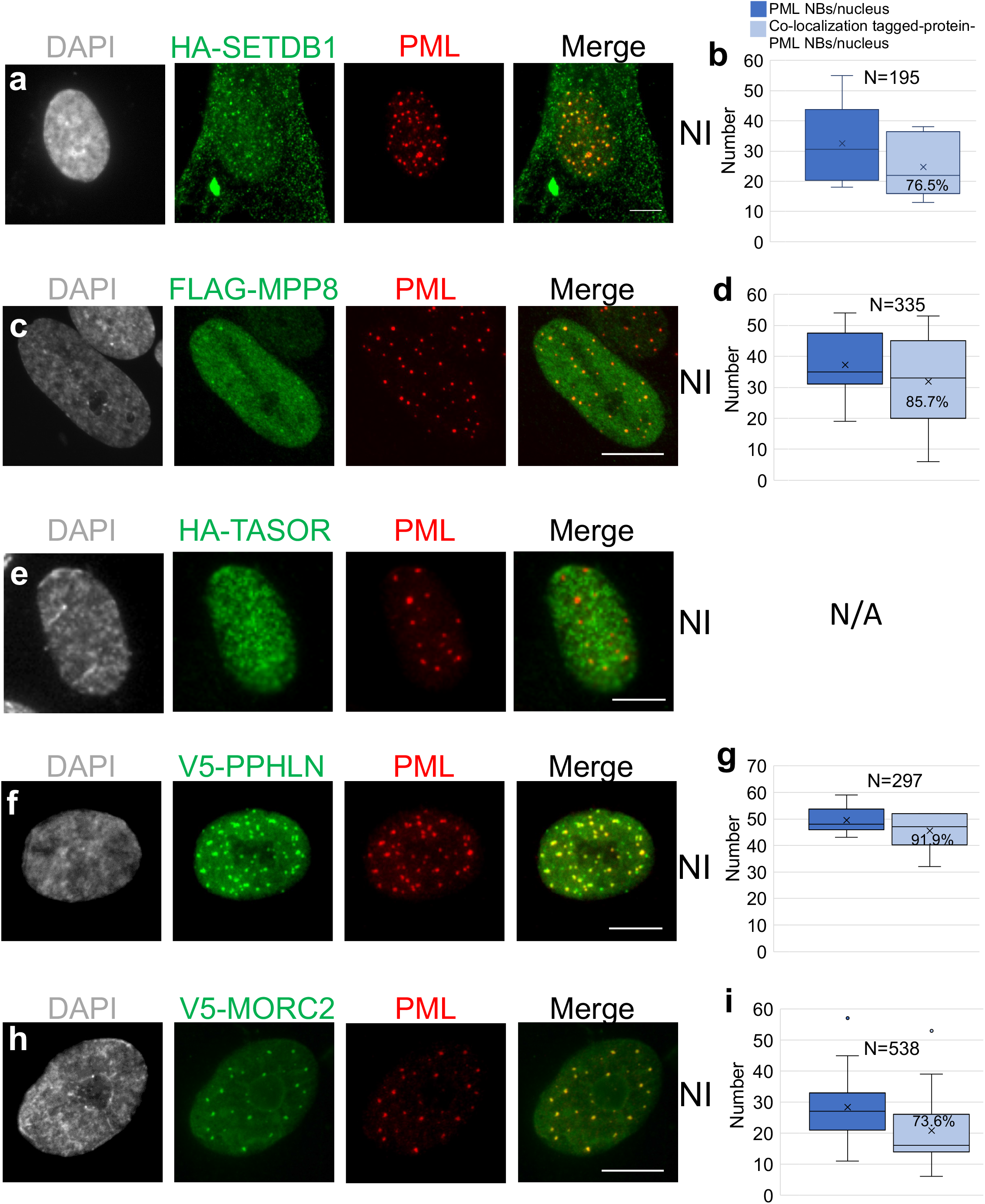
Ectopically expressed SETDB1, MPP8, PPHLN1, MORC2 co-localize with PML NBs. **a, c, e, f, h** Immunofluorescence on hFC transfected with plasmids expressing tagged versions of proteins of interest (green) co-stained with PML (red). Scale bars represent 5µm. **b, d, g, i** quantification of the average number of PML NBs per nucleus (dark blue) and of the co-localization of the protein with the PML NBs (light blue) for the respective samples in non-infected (NI) cells. N = number of events counted. Means are indicated in the graphs.

**Supplementary Fig.4.**
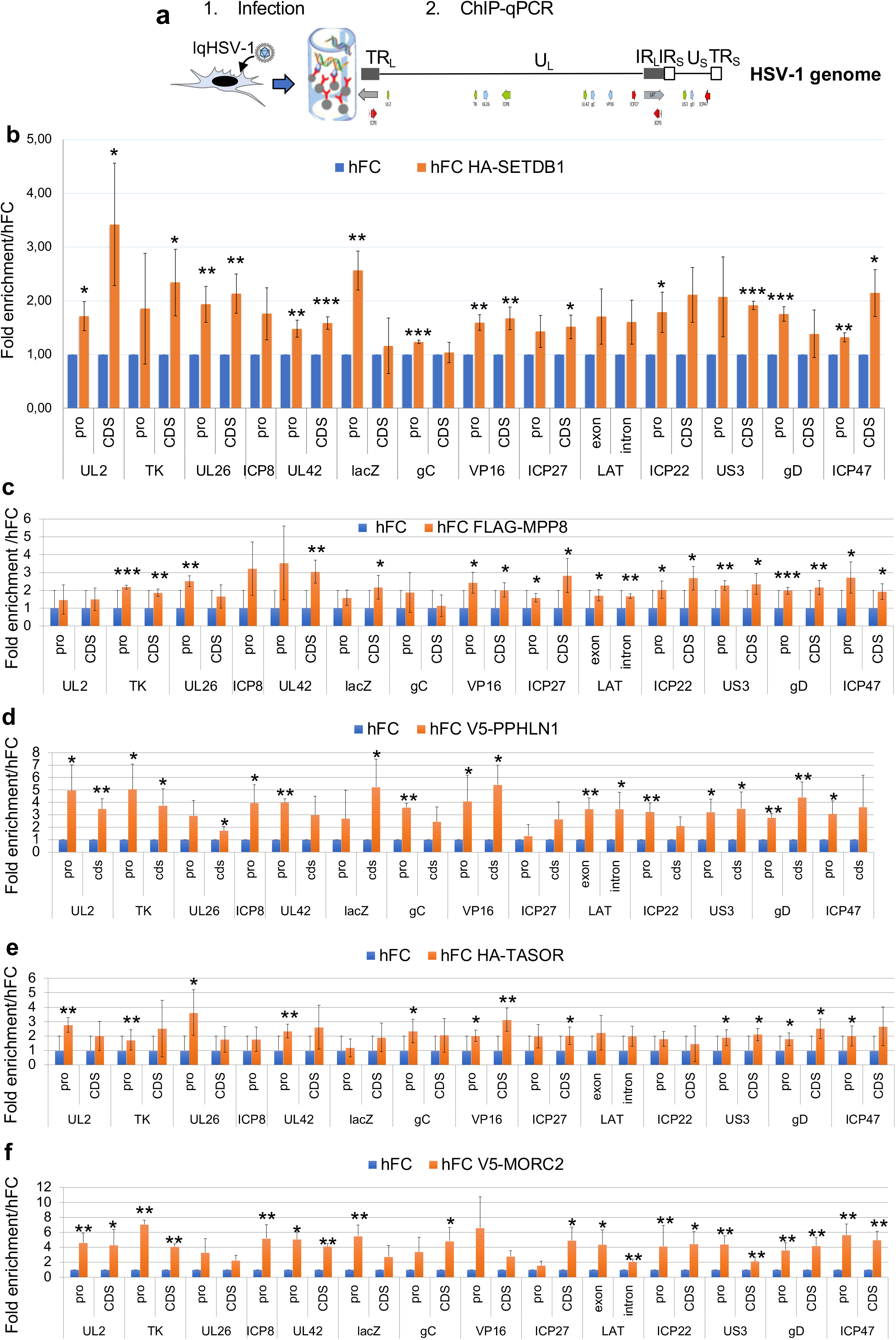
Expanded ChIP-qPCR data for the interaction of proteins of the HUSH-SETDB1-MORC2 entity with PML NBs-associated quiescent HSV-1. **a,** schematic view of the ChIP procedure to detect interactions of proteins with viral genomes. **b to f,** control hFC or hFC expressing a tagged version of proteins of interest were infected with lqHSV-1 (m.o.i.= 3) and ChIP-qPCR was performed 24h pi. All experiments were performed at least 3 times. Measurements were taken from distinct samples. P-values * <0.05; ** <0.01 (one-tailed paired Student’s t-test).

**Supplementary Fig. 5.**
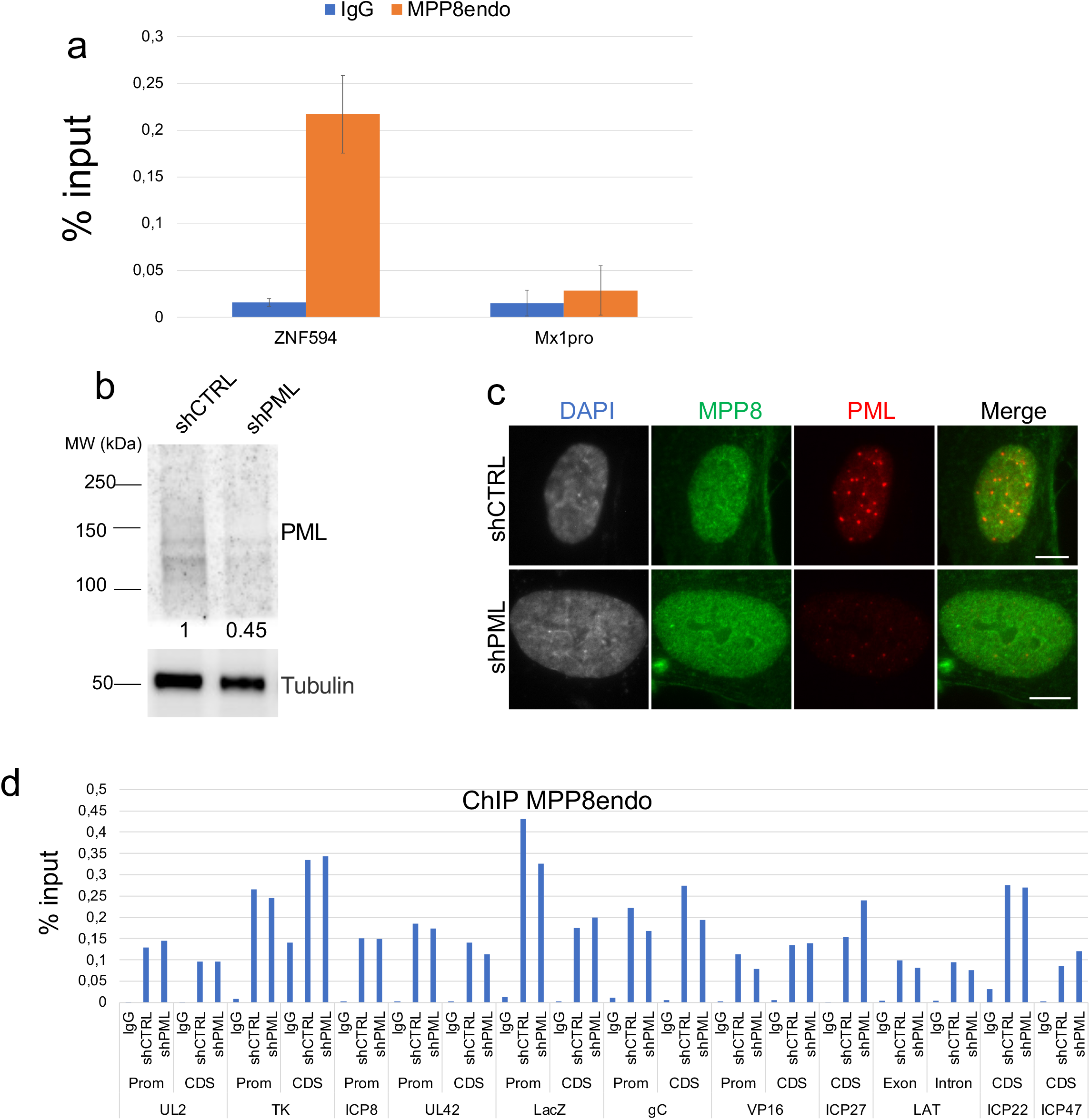
Depletion of PML does not impact on binding of endogenous MPP8 with lqHSV-1. **a,** Interaction of endogenous MPP8 with cellular loci. hFC were infected with lqHSV-1 (m.o.i.= 3) and ChIP-qPCR was performed 24h pi. The data show the interaction of endogenous MPP8 with a representative ZNF594 cellular locus according to (27) as well Mx1 promoter (Mx1pro) not known to bind HUSH (mean values +/- SD). Data from 2 independent experiments. b, Western blotting of PML in shCTRL or shPML-expressing hFC. Tubulin is shown as loading control. The quantification of PML relative to tubulin is depicted below the WB. c, immunofluorescence showing the PML NBs (red), MPP8 (green), and nuclei (grey) signals in shCTRL or shPML-expressing hFC. Bar represents 5µm. d, shCTRL or shPML-expressing hFC were infected with lqHSV-1 (m.o.i.= 3) and ChIP-qPCR was performed 24h pi. ChIP-qPCR was performed using MPP8 antibody and IgG as control antibody to detect MPP8 interaction with viral genome loci.

**Supplementary Fig. 6.**
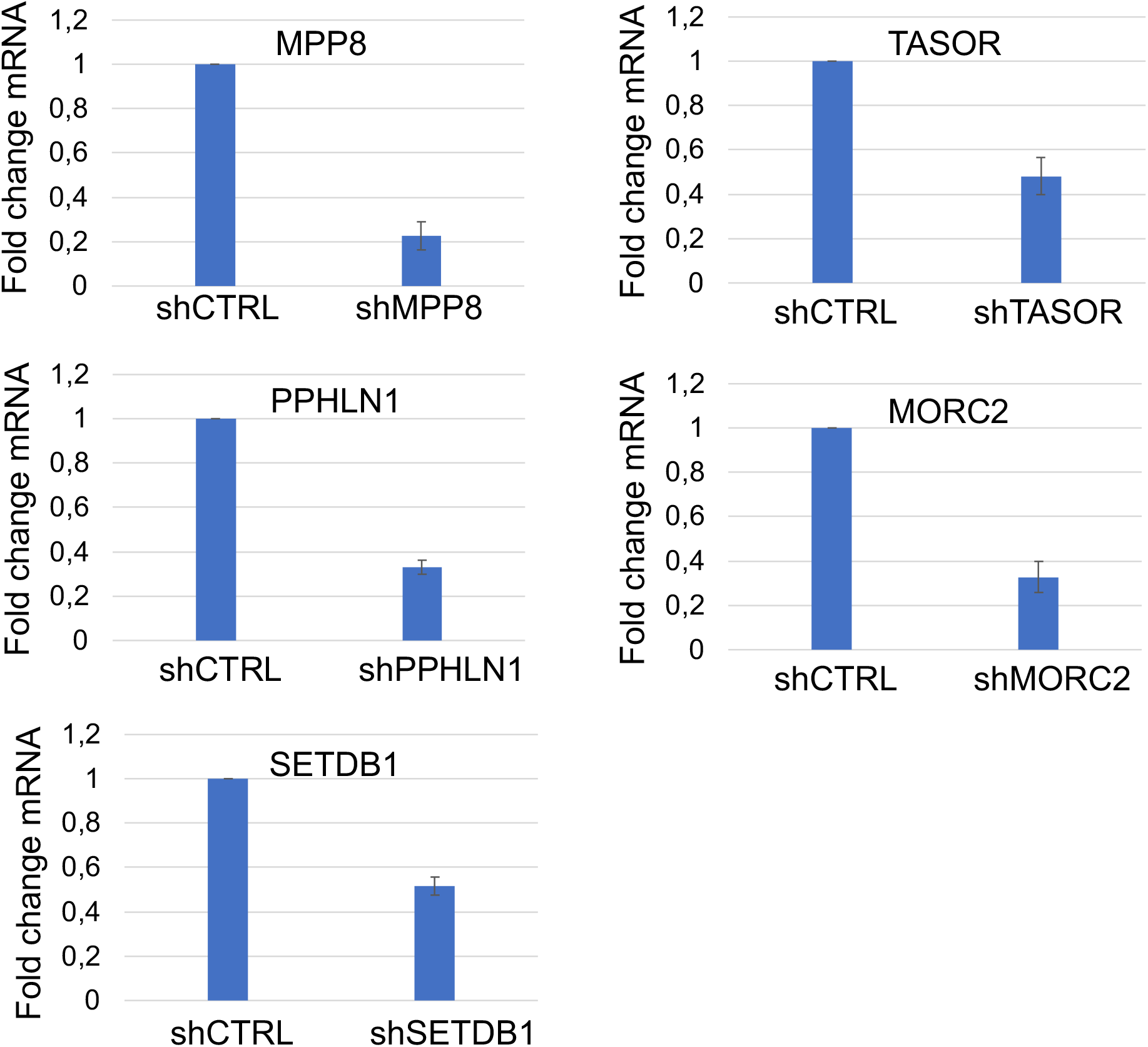
Quantifications of mRNA of HUSH-SETDB1-MORC2 in shRNA-treated hFC. a, hFC were transduced with lentiviruses expressing control shRNA (shCTRL) or shRNA against proteins before infection with lqHSV-1 for 48h (m.o.i.= 3). Cells were harvested and RT-qPCR was performed.

**Supplementary Fig. 7.**
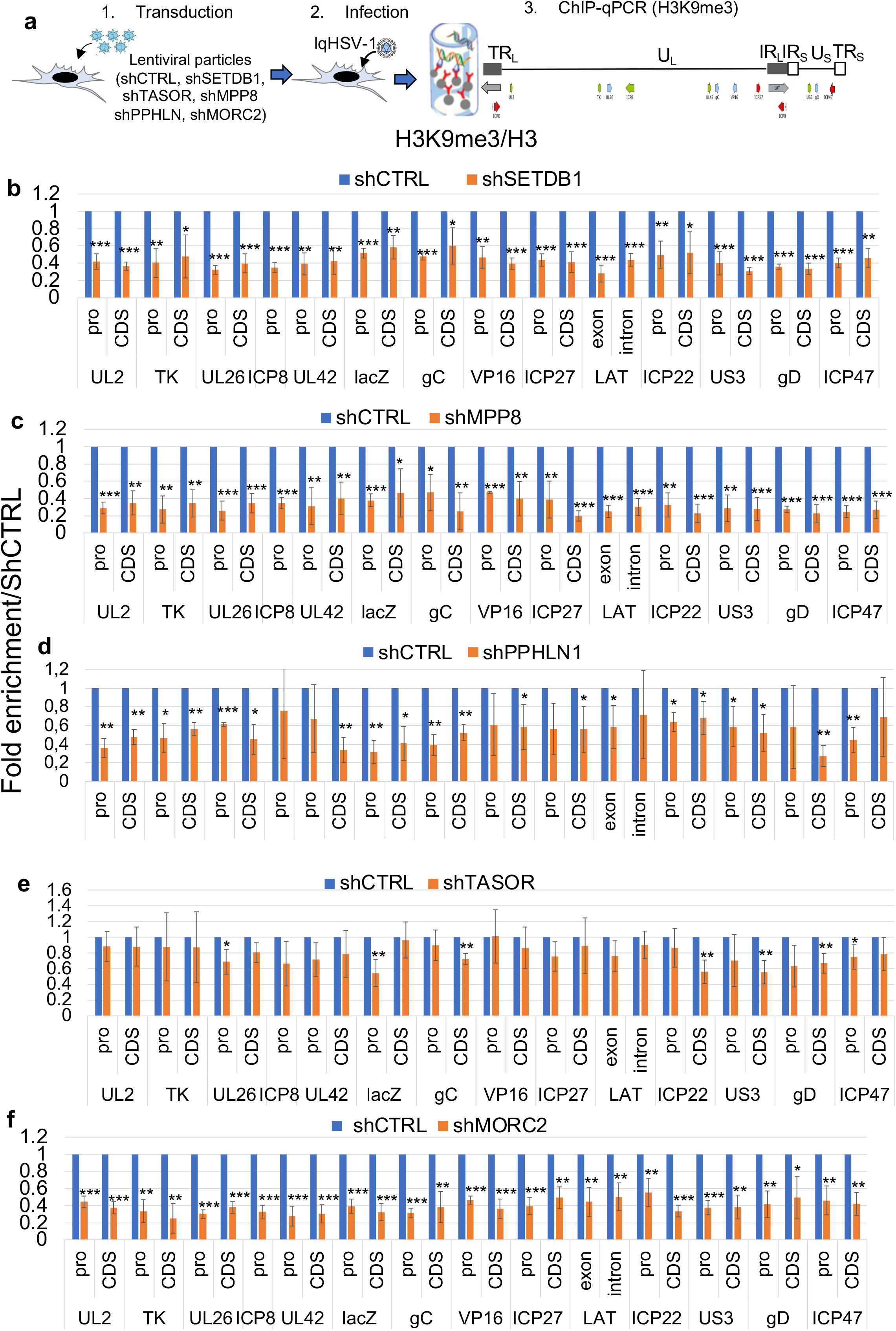
Expanded ChIP-qPCR data for the impact of depletion of the proteins of the HUSH-SETDB1-MORC2 entity on H3K9me3 association with lqHSV-1. **a,** schematic view of the ChIP procedure to detect interactions of H3K9me3 with viral genomes after depletion or not by shRNAs of proteins of interest. **b to f,** hFC expressing a shRNA control or hFC expressing a shRNA inactivating a specific protein were infected with lqHSV-1 (m.o.i.= 3) and ChIP-qPCR was performed 24h pi. H3K9me3 signal was standardized over total H3 present on each viral locus. All experiments were performed at least 3 times. Measurements were taken from distinct samples. P-values * <0.05; ** <0.01 (one-tailed paired Student’s t-test).

**Supplementary Fig. 8.**
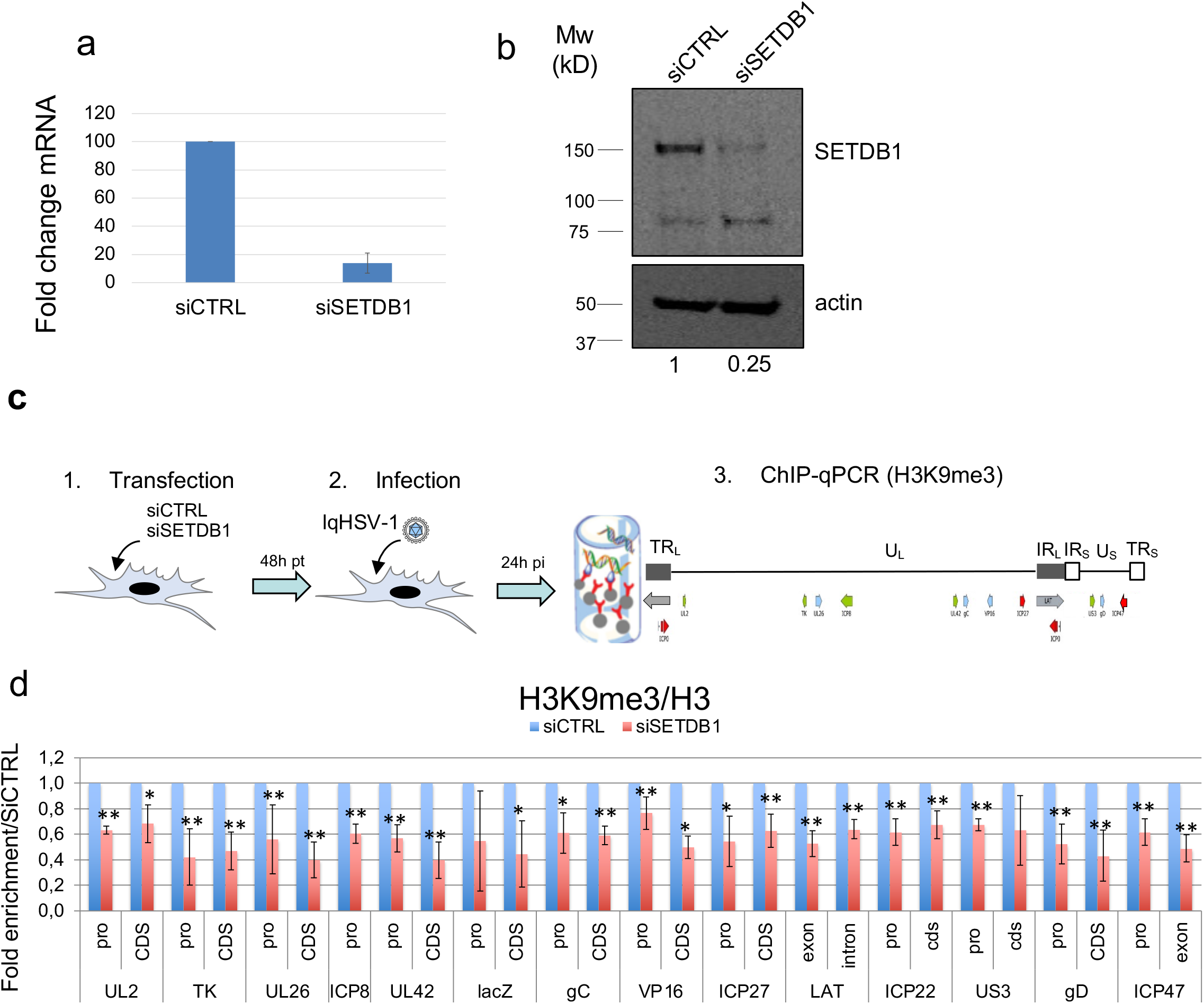
Downregulation of SETDB1 with a siRNA also impacts on H3K9me3 association with lqHSV-1. **a,** RT-qPCR showing the efficiency of the siRNA SETDB1 to reduce SETDB1 mRNA. **b,** WB analysis showing that SETDB1 is impacted at the protein level by the use of the siRNA SETDB1. The quantification of SETDB1 relative to actin is depicted below the WB. **c,** schematic view of the ChIP procedure to detect interactions of H3K9me3 with viral genomes after depletion or not by siRNAs of SETDB1. **d,** hFC expressing a siRNA control (luciferase) or hFC expressing a siRNA inactivating SETDB1 were infected with lqHSV-1 (m.o.i.= 3) and ChIP-qPCR was performed 24h pi. H3K9me3 signal was standardized over total H3 present on each viral locus. Experiments were performed at least 3 times. Measurements were taken from distinct samples. P-values * <0.05; ** <0.01 (one-tailed paired Student’s t-test).

**Supplementary Fig. 9.**
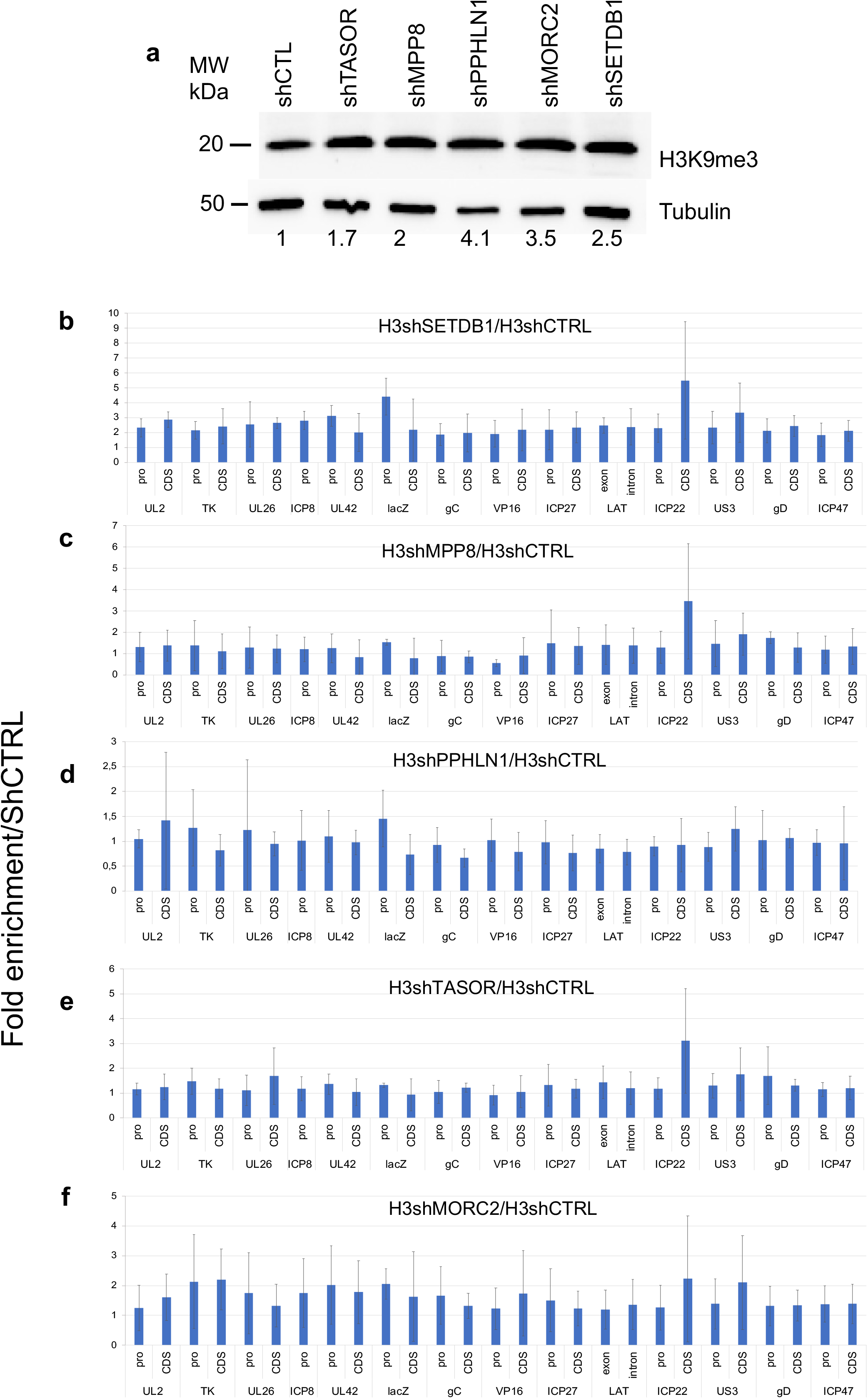
a, HUSH-SETDB1-MORC2 depletion does not reduce the total level of H3K9me3. **a,** Detection of total H3K9me3 in cells depleted of the HUSH-SETDB1-MORC2 by shRNAs. The quantification of H3K9me3 relative to tubulin is depicted below the WB. **b to f. HUSH-SETDB1-MORC2 depletion does not impact on histone H3 association with lqHSV-1.** hFC expressing a shRNA control or hFC expressing a shRNA inactivating a specific protein were infected with lqHSV-1 (m.o.i.= 3) and ChIP-qPCR was performed 24h pi. Total histone H3 present on each viral locus was measured comparing the specific shRNA to the shCTRL conditions. All experiments were performed at least 3 times.

**Supplementary Fig. 10.**
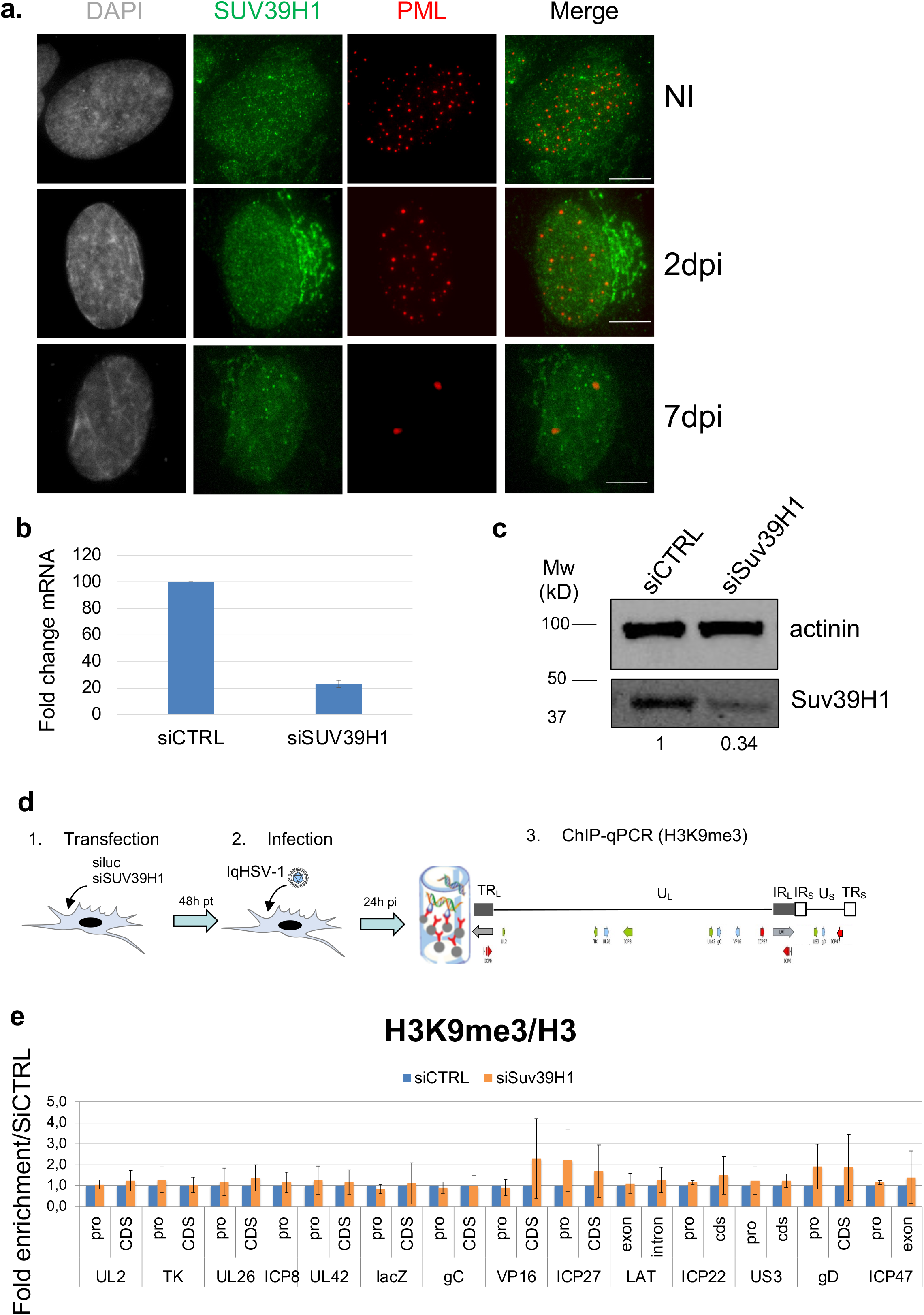
SUV39H1 is not involved in H3K9me3 association with lqHSV-1. **a,** hFC were infected or not (NI) with lqHSV-1 (m.o.i. 3, 100% of cells infected) and harvested at different times post-infection (2, 7 dpi) to perform immunofluorescence to detect SUV39H1 (green), PML NBs (red), and nuclei (DAPI, grey). Scale bar represents 5µm. **b,** RT-qPCR showing the efficiency of the siRNA SUV39H1 to reduce SUV39H1 mRNA. **c,** WB analysis showing that SUV39H1 is impacted at the protein level by the use of the siRNA SUV39H1. The quantification of SUV39H1 relative to actinin is depicted below the WB. **d,** schematic view of the ChIP procedure to detect interactions of H3K9me3 with viral genomes after depletion or not by siRNAs of SUV39H1. **e,** hFC expressing a siRNA control (luciferase) or hFC expressing a siRNA inactivating SUV39H1 were infected with lqHSV-1 (m.o.i.= 3) and ChIP-qPCR was performed 24h pi. H3K9me3 signal was standardized over total H3 present on each viral locus. Experiments were performed at least 3 times. Measurements were taken from distinct samples.

**Supplementary Fig. 11.**
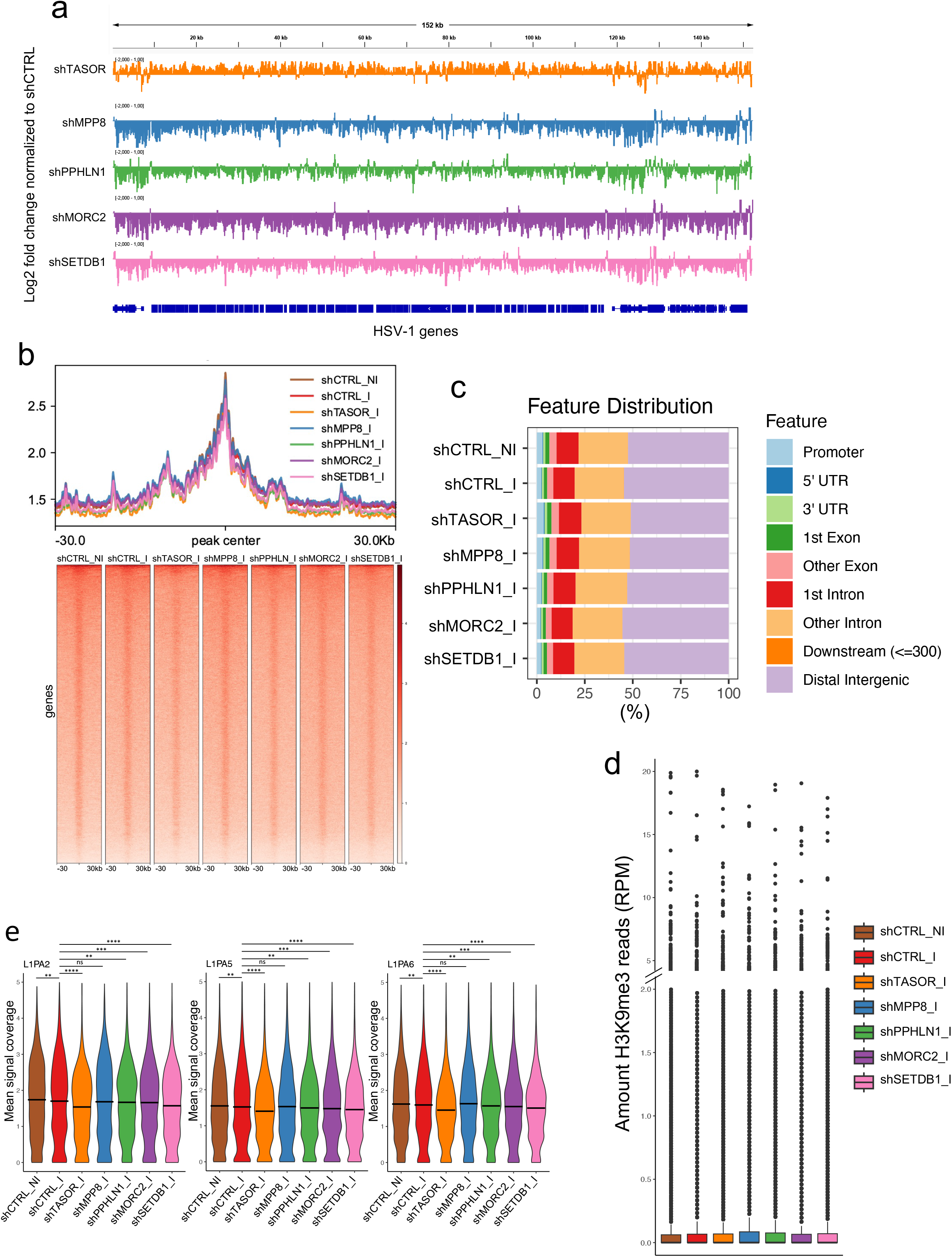
ChIP-seq data analyses for the impact of depletion of the proteins of the HUSH-MORC2 entity on H3K9me3 association with lqHSV-1 and cellular genes. Data from H3K9me3 ChIP-seq experiments in infected hFC treated with shCTRL or a shRNA against the indicated proteins of interest. **a,** Genome browser snapshot of the H3K9me3 enrichment, normalized as a log2 fold change to shCTRL, across the entire PML NBs-associated quiescent HSV-1 genome. **b,** Profile plot and heatmaps showing the density of H3K9me3 signal on large H3K9me3-rich regions defined in (42). **c,** Genomic feature distribution of identified peaks. **d,** Box plots showing the quantification of H3K9me3 reads on the cellular genes (in Reads Per Million). **e**, ChIP-Seq mean signal coverage of H3K9me3 on different LINEs subtypes across the different samples (related to figures 3m, n, o). Y-axis was adjusted to a 0-5 range. Adjusted p-values ** <0.01; *** <0.001; *** <0.001 (Wilcoxon rank-sum test), ns: non significant.

**Supplementary Fig. 12.**
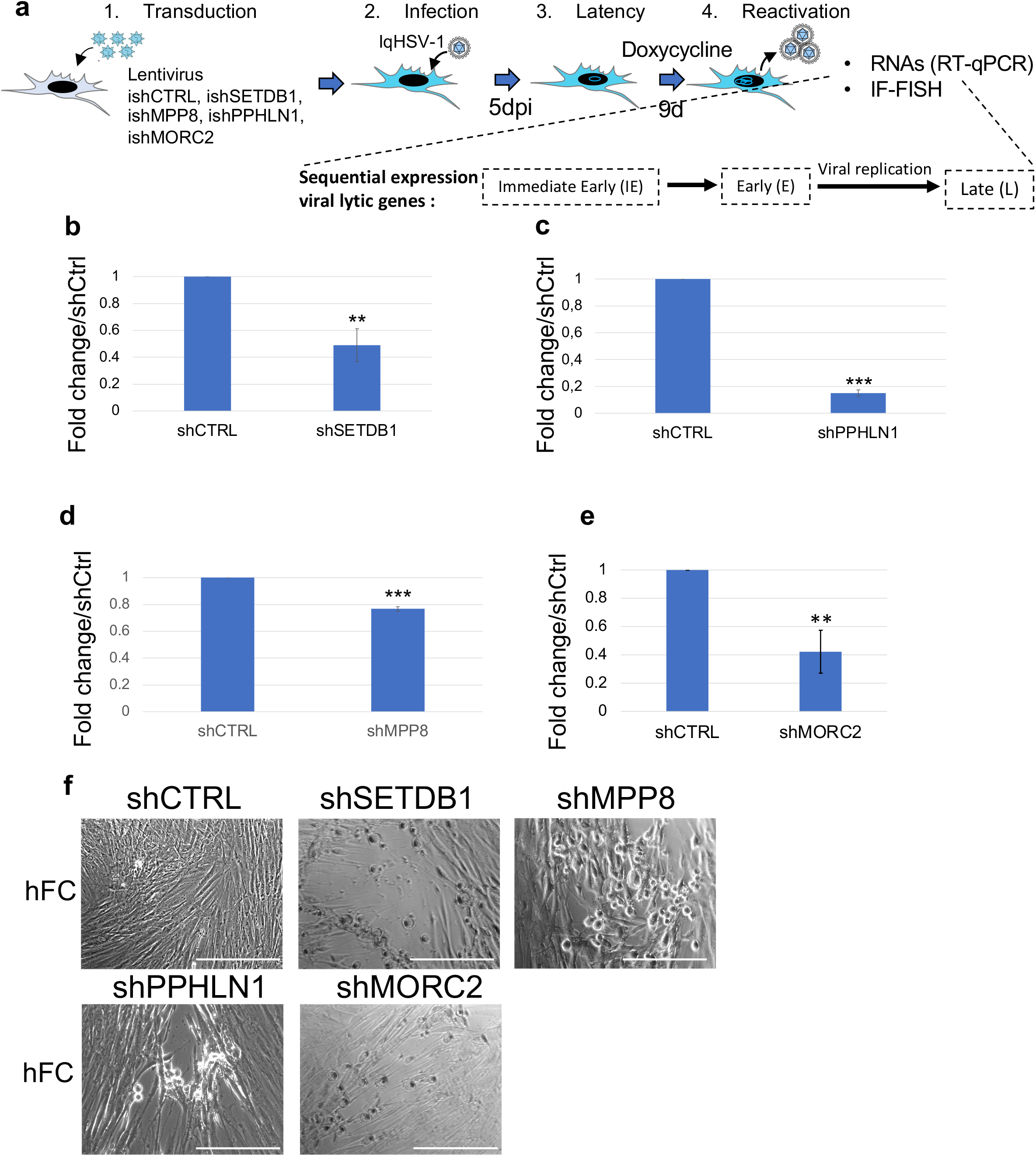
HUSH-SETDB1-MORC2 mRNAs quantification following inducible shRNAs expression. **a,** schematic view of the experimental procedure to determine the restrictive activity of the individual proteins of the HUSH-SETDB1-MORC2 entity. **b-e**, RT-qPCR performed from dox inducible shRNAs expressing cells to quantify SETDB1, PPHLN1, MPP8, MORC2 mRNA, respectively. All experiments were performed at least 3 times. Measurements were taken from distinct samples. All quantifications data are mean values (+/- SD). P-values * <0.05; ** <0.01; *** <0.001 (one-tailed paired Student’s t-test). **f,** representative images of hFC monolayers showing viral plaques appearing as a result of virus reactivation and spread, following downregulation of individual proteins by shRNA. Scale bars represent 100 µm.

**Supplementary Fig. 13.**
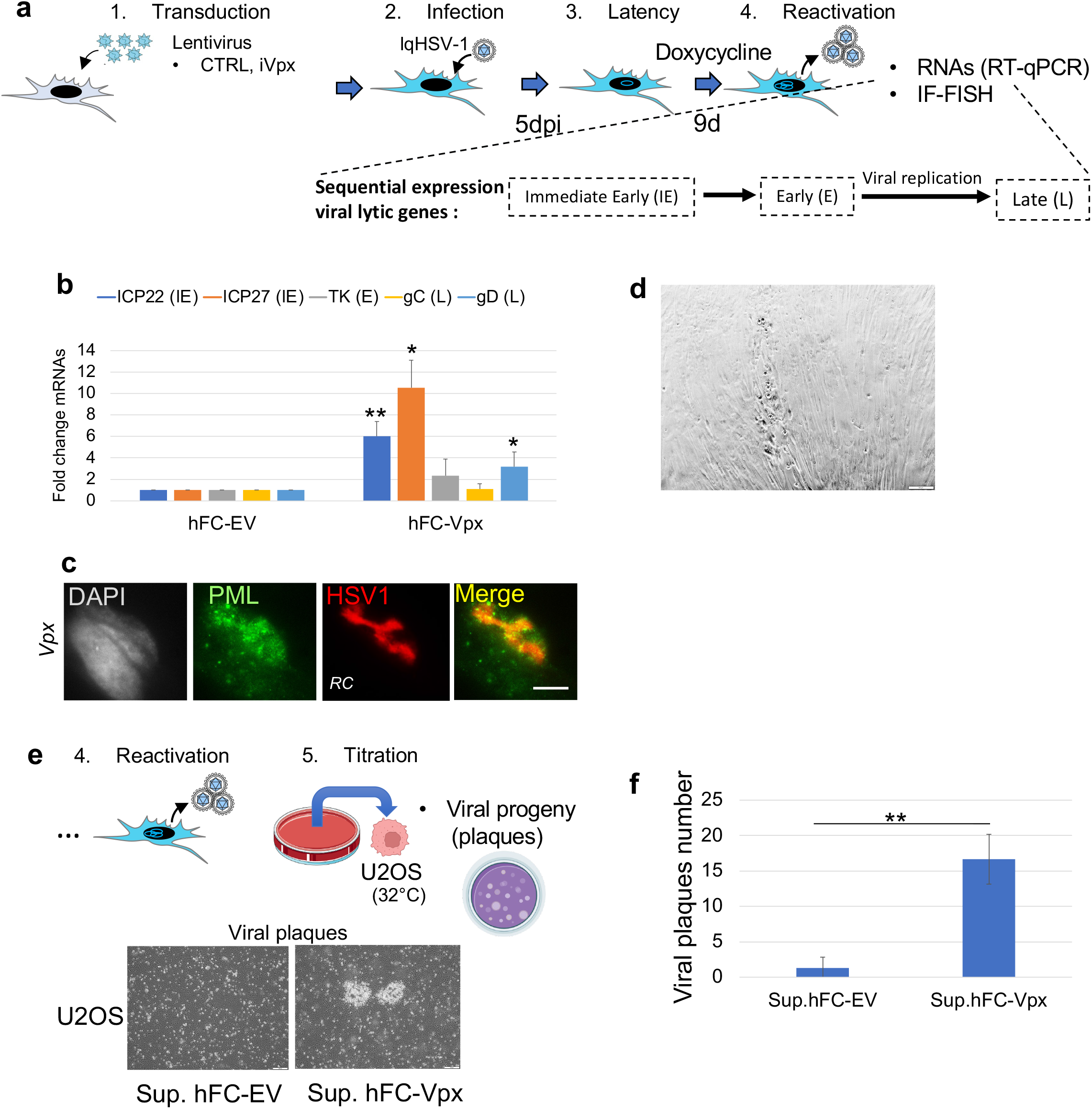
Expression of HIV-2_ROD_ Vpx induces HSV-1 reactivation. **a,** schematic view of the experimental procedure as described in figure 3. **b,** RT-qPCR on representative HSV-1 lytic genes following doxycycline treatment of hFC transduced with an empty pLVX vector (hFC-EV) or a pLVX vector expressing Vpx HIV-2_ROD_ (hFC-Vpx). **c,** hFC monolayer showing a viral plaque appearing following Vpx HIV-2_ROD_ expression. Scale bar represents 100 µm. **d,** immuno-FISH allowing the detection of PML (green), HSV-1 (red) and nuclei (grey, DAPI) following Vpx HIV-2_ROD_ expression. Scale bar represents 5 µm. RC: replication compartment**. e,** titration on U2OS cells of infectious viral progeny production as described in figure 3. Up: procedure; down: viral plaques with supernatant from Vpx HIV-2_ROD_-expressing hFC. **f,** numeration on U2OS cells of infectious viral progeny production in control hFC vs Vpx HIV-2_ROD_-expressing hFC. Scale bar represents 100 µm.

**Supplementary Fig. 14.**
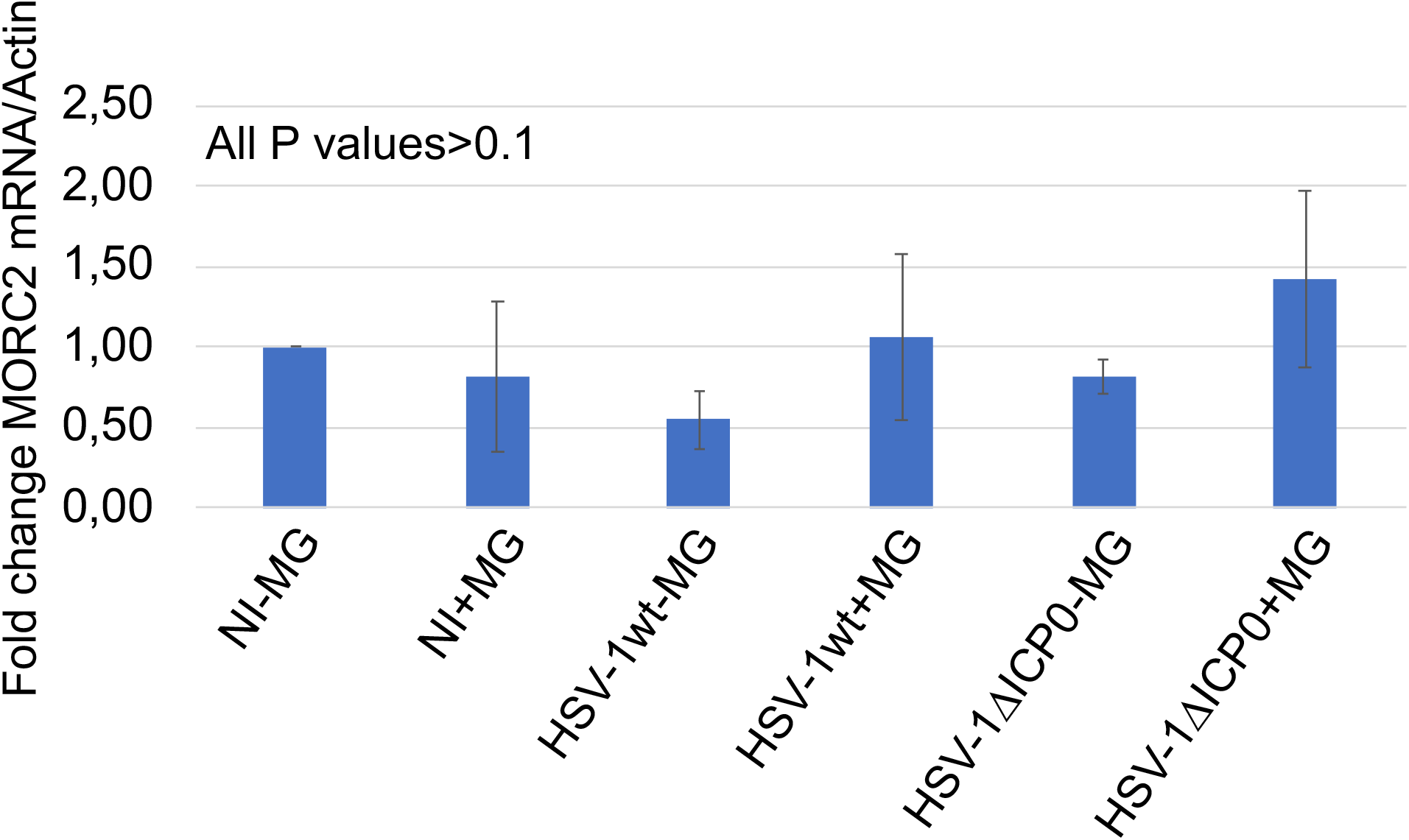
Quantification of MORC2 mRNA in infected cells. hFC were infected or not with wild type HSV-1 (HSV-1wt) or a mutant HSV-1 unable to express ICP0 (HSV-1ΔICP0), in the presence (+) or not (-) of the proteasome inhibitor MG132 (MG). Eight hours post infection cells were treated to extract total RNA and perform RT-qPCR on MORC2 mRNA. Actin mRNA was used as control to adjust mRNA quantity from samples to samples.

**Supplementary Fig. 15.**
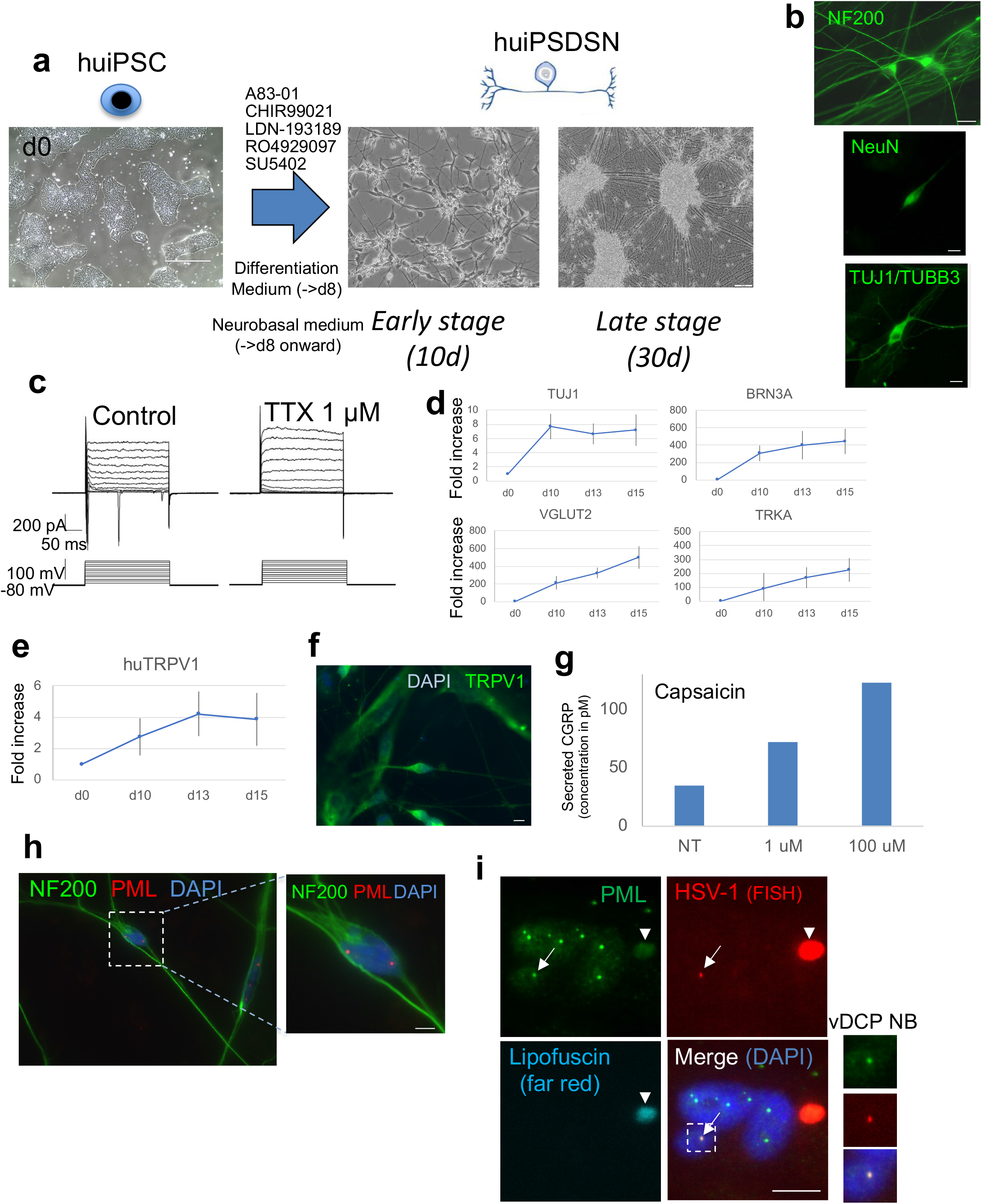
hiPSDN have nociceptive neurons characteristics and contain PML NBs. **a,** Phase contrast images of cultured hiPSC and resulting hiPSDN after differentiation protocol described in the methods section. Scale bar represents 10 µm. **b,** immunofluorescence staining of hiPSDN for three major neuronal markers intermediate neurofilament (NF200), neuronal nuclear antigen (NeuN), and beta-III Tubulin (TUJ1/TUBB3). Scale bar represents 10 µm. **c,** Current-clamp recording of membrane potential in iPSC-derived neurons: example recording of a single action potential evoked by current injection of 200 pA for 50 ms. Application of TTX blocked the generation of action potentials. **d,** Representative RT-qPCR (N=3) to measure the relative fold increase in the expression of neuronal (TUJ1/TUBB3) and sensory (BRN3A, vGLUT2, TRKA) neurons from d0 (hiPSC) to d15 (hiPSDN). **e,** RT-qPCR (N=3) to measure the relative fold increase over time in the expression of the TRPV1 receptor, a marker of nociceptive neurons. **f,** immunofluorescence staining of the TRPV1 receptor in hiPSDN. Scale bar represents 10 µm. **g,** functional analysis of nociceptive hiPSDN for their capacity to respond to capsaicin stimulus by the secretion of CGRP peptide. **h,** immunofluorescence staining showing the presence of native PML NBs (red) in hiPSDN (NF200, green). **i,** FISH-IF detection of HSV-1 genome (red) associated to PML NBs (green) (vDCP NBs, arrows). Lipofuscin (arrow heads) is an autofluorescent cytoplasmic signal detected in human neurons and recognizable by its fluorescence in all the channels including far red (already described in (4)). Scale bar represents 10 µm.

**Supplementary Fig. 16.**
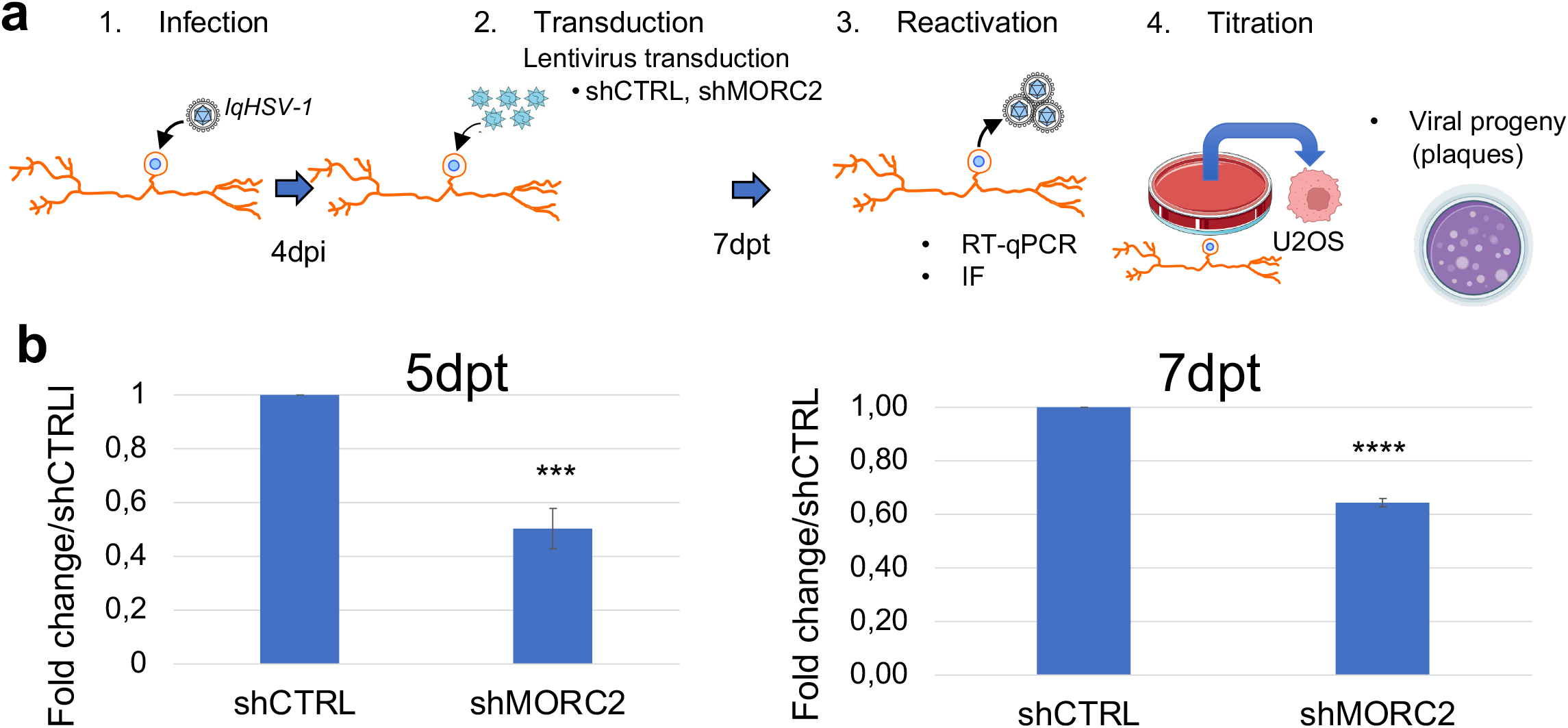
MORC2 shRNA efficiently reduces MORC2 mRNA in iPSDN. **a,** schematic view of the experimental procedure as described in figure 6. **b**, RT-qPCR at 5dpt (left) and 7dpt (right) for the quantification of MORC2 mRNAs. Experiments are in triplicates. Measurements were taken from distinct samples. Quantifications data are mean values (+/- SD). P-values *** <0.001 (one-tailed paired Student’s t-test).

